# Integrated data from R405W desmin knock-in mice highlight alterations of mitochondrial function, protein quality control, and myofibrillar structure in the initial stages of myofibrillar myopathy

**DOI:** 10.1101/2024.09.29.615655

**Authors:** Sabrina Batonnet-Pichon, Florence Delort, Alain Lilienbaum, Carolin Berwanger, Dorothea Schultheis, Ursula Schlötzer-Schrehardt, Andreas Schmidt, Steffen Uebe, Yosra Baiche, Tom J. Eisenack, Débora Broch Trentini, Markus Mallek, Leonid Mill, Ana Ferreiro, Bettina Eberhard, Thomas Lücke, Markus Krüger, Christian Thiel, Rolf Schröder, Christoph S. Clemen

**Author notes:** Authors for correspondence: Sabrina Batonnet-Pichon, Institut Cochin, Université Paris Cité, INSERM U1016, CNRS, F-75014 Paris, France, 75014 Paris, France, Rolf Schröder, Institute of Neuropathology, University Hospital Erlangen, Schwabachanlage 6, 91054 Erlangen, Germany, Christoph S. Clemen, Institute of Aerospace Medicine, German Aerospace Center (DLR), Linder Höhe, 51147 Cologne, Germany. These three first authors contributed equally to this work (Sabrina Batonnet-Pichon, Florence Delort, Alain Lilienbaum). These two senior authors contributed equally to this work (Rolf Schröder, Christoph S. Clemen).

## Abstract

**Background:** Mutations in the desmin gene cause skeletal myopathies and cardiomyopathies. The objective of this study was to elucidate the molecular pathology induced by the expression of R405W mutant desmin in murine skeletal muscle tissue.

**Methods:** A comprehensive characterization of the skeletal muscle pathology in hetero- and homozygous R405W desmin knock-in mice was performed, employing grip strength, blood acylcarnitine and amino acid, histological, ultrastructural, immunofluorescence, immunoblot, ribosomal stalling, RNA sequencing and proteomic analyses.

**Results:** Both hetero- and homozygous R405W desmin knock-in mice showed classical myopathological features of a myofibrillar myopathy with desmin-positive protein aggregation, degenerative changes of the myofibrillar apparatus, increased autophagic build-up, and mitochondrial alterations. Muscle weakness and increased blood concentrations of acylcarnitines and amino acids were only present in homozygous animals. During its translation, mutant desmin does not induce terminal ribosomal stalling. Analyses of RNA sequencing and proteomic data from soleus muscle of 3-month-old mice depicted 59 up- and 2 down-regulated mRNAs and 101 up- and 18 down-regulated proteins that were shared between the heterozygous and homozygous genotypes in the respective omics datasets compared to the wild-type genotype. Combined analysis of the omics data demonstrated 187 significantly dysregulated candidates distributed across four groups of regulation. A down-regulation on the mRNA and protein levels was observed for a multitude of mitochondrial proteins including essential proton gradient-dependent carriers. Up-regulation on both omics levels was present for the transcription factor Mlf1, which is a binding partner of protein quality control related Dnajb6. Down-regulated on mRNA but up-regulated on the protein level was the sarcomeric lesion marker Xirp2 (xin actin-binding repeat-containing protein 2), whereas Ces2c (acylcarnitine hydrolase) was regulated in the opposite way.

**Conclusions:** The present study demonstrates that the expression of mutant desmin results in a myofibrillar myopathy in hetero- and homozygous R405W desmin knock-in mice. Combined morphological, transcriptomic and proteomic analyses helped to decipher the complex pattern of early pathological changes induced by the expression of mutant desmin. Our findings highlight the importance of major mitochondrial alterations, including essential proton gradient-dependent carriers as well as Dnajb6-related protein quality control and Xin-related myofibrillar damage, in the molecular pathogenesis of desminopathies.

## Introduction

Desmin, a muscle specific member of the large family of intermediate filament proteins, is a key component of the extrasarcomeric cytoskeleton in muscle cells [1]. This three-dimensional filamentous cytoskeletal structure organizes the proper alignment of neighboring myofibrils at the level of Z-discs [2], interlinks and attaches the myofibrillar apparatus to costameres in skeletal muscle [3–5], to intercalated discs in cardiac cells [6, 7], and to myotendinous junctions [8] and neuromuscular junctions [9] as well as mitochondria [10–12] and nuclei [13–15] in striated muscle cells. Beyond these roles in the spatial organization of myofibrils and the positioning of cell organelles, desmin has been reported to exert multiple roles in mechanosensing and stress tolerance [16, 17], adhesion and migration [18], and cell signaling [19]. The essential role of desmin in human skeletal muscle is highlighted by the observation that mutations in the human desmin gene (*DES*) cause autosomal-dominant, autosomal-recessive and sporadic myopathies and cardiomyopathies [2, 20]. In the last three decades more than 100 disease-causing human *DES* mutations have been described [2, 21]. The term ‘desminopathies’ comprises all *DES* mutation-related disorders irrespective of their form of inheritance, mutation type and their consequences on desmin protein expression. Autosomal-dominant forms, by far the most frequent form of human desminopathies, typically display desmin-positive protein aggregation pathology and myofibrillar disarray in striated muscle tissue [2]. In the very rare and clinically more severe autosomal-recessive cases due to homozygous or compound heterozygous *DES* mutations, delineated a pronounced cardiotoxic effect of the issue of desmin and desmin protein aggregation is more complex. In one subform the desmin protein expression is completely abolished and there is no protein aggregation pathology [22–25], whereas in a second and third subform reduced protein levels of solely mutant desmin are seen in conjunction with [26–36] or without [37] concomitant signs of protein aggregation. The desminopathies with protein aggregation in striated muscle tissue belong to the clinically and genetically heterogeneous group of myofibrillar myopathies [38], which is a numerically significant subgroup of human protein aggregation myopathies [39]. The precise mechanisms leading from the genetic defect to the histopathological lesions observed are poorly understood and, to date, no specific treatment is available for desminopathies and all other forms of myofibrillar myopathies [2].

We previously reported on the pathogenic effects of the human R406W desmin missense mutation in a young male patient with progressive restrictive cardiomyopathy, cardiac conduction defects, and arrhythmias necessitating heart transplantation as well as on the generation and characterization of hetero- and homozygous R405W desmin knock-in mice, which harbor the ortholog of the human R406W desmin mutation [40]. In addition to previously reported patients harboring this particular missense mutation, who all showed an early disease onset and severe cardiac disease [27, 41–45], our work in humans and mice delineated a pronounced cardiotoxic effect of R406W/R405W desmin based on a compromised attachment of structurally altered desmin filaments to intercalated discs [40]. To further decipher the pathology and molecular mechanisms inflicted by mutant desmin, we report here on the comprehensive characterization of the skeletal muscle pathology in hetero- and homozygous R405W desmin knock-in mice by means of grip strength, histological, ultrastructural, immunofluorescence, immunoblot, ribosomal stalling, RNA sequencing, and proteomic analyses.

## Materials and Methods

### R405W desmin knock-in mice

The R405W desmin knock-in mouse model C57BL/6N-*Des*^tm1.1Allb^ (http://www.informatics.jax.org/allele/MGI:6382607) was generated as described previously [40]. Routine genotyping on gDNA from tail biopsies was performed by PCR using the primer pair 5’-CTGGAGGAGGAGATCCGACA-3’ and 5’-GGCCCTCGTTAATTTTCTGC-3’. Mice were housed in isolated ventilated cages under specific and opportunistic pathogen-free conditions at a standard environment with free access to water and food. Muscle strength was tested using a grip strength meter (Bioseb BIO-GS3+, Bioseb, Vitrolles, France). Health monitoring was done as recommended by the Federation of European Laboratory Animal Science Associations. Mice were handled in accordance with European Union guidelines and French regulations for animal experimentation, and the investigations were approved by the University Paris Diderot local committee (authorization number CEB-16-2016 / 2016041216476300). Moreover, the Buffon animal facility (Plateforme d’Hébergement et d’expérimentation animale Buffon) is fully licensed by French competent authorities and has animal welfare insurance.

### Histological evaluation

Murine soleus/gastrocnemius muscle packages were dissected and immediately embedded in Tissue-Tek OCT Compound (Sakura Finetek, Torrance, CA, USA), frozen in liquid nitrogen-cooled isopentane, and stored at −80°C. Cryostat sections of 6 µm thickness were collected on microscope slides, air-dried for 30 min, and used for standard histology staining and evaluation. In addition, a deep learning-based image analysis approach using photo-realistic computer graphics generated synthetic data in muscle histopathology [46] was used to quantitatively analyse the soleus muscle part in hematoxylin and eosin stained sections derived from 3-month-old mice.

### Immunoblotting and antibodies

For the extraction of proteins, snap-frozen soleus muscles were pulverized in a mortar on liquid nitrogen before addition of lysis buffer (50 mM TrisHCl pH 8.0, NaCl 150 mM, EDTA 1 mM, NP40 1%). Samples were rotated at 4°C for 1h and then sonicated for 10 s. Samples were not centrifugated to avoid loss of desmin, used undiluted for protein quantitation by the Bradford method, and then 1:5 diluted with 1x SDS sample buffer (25 mM Tris, 0.8% SDS, 2% 2-mercaptoethanol, 4% glycerol, 0.001% bromophenol blue, pH 6.8) and boiled at 95°C for 7 min. A total of 30 µg of muscle extracts were loaded per line on a 10 or 15% acrylamide gel for SDS-PAGE. After separation, proteins were transferred onto nitrocellulose membranes (0.45 µm, Macherey Nagel), which were blocked 2h with either 5% non-fat milk or 4% BSA with low endotoxin, IgG and proteases (US Biological) in 0.5% Tween/PBS solution. Membranes were incubated with polyclonal rabbit anti-p62 (SQSTM1) (P0067, Sigma-Aldrich, 1:500) for 1h at room temperature. Anti-rabbit secondary antibody coupled with horseradish peroxidase (#31460, Pierce/Thermo Scientific, 1:10,000) was detected following incubation with Clarity Western ECL (BioRad) and visualized with a CCD camera (FUJI LAS 4000, GE Healthcare).

### Mass spectrometric analysis of acylcarnitine and amino acid levels in blood

Mass spectrometric quantitation of acylcarnitines and amino acids from murine retro-orbital sinus blood samples was performed as previously described in detail [47].

### Electron microscopy

Soleus muscle specimens were fixed in 2.5% glutaraldehyde in 0.1 M phosphate buffer, pH 7.2, post-fixed in 2% buffered osmium tetroxide, dehydrated in graded alcohol concentrations, and embedded in epoxy resin according to standard protocols. 1 µm semi-thin sections for orientation were stained with toluidine blue. Ultra-thin sections were stained with uranyl acetate and lead citrate, and examined with a LEO 906E transmission electron microscope (Carl Zeiss GmbH, Oberkochen, Germany) at the Department of Ophthalmology, University Hospital Erlangen, Friedrich-Alexander University Erlangen-Nürnberg, Erlangen, Germany, or a JEOL 1011 transmission electron microscope (JEOL GmbH, Freising, Germany) at the Electron Microscopy Platform, Cochin Institute, Paris, France.

### Immunofluorescence microscopy and antibodies

Muscle tissue sections were fixed for 10 min with acetone at −20°C, and air-dried for 10 min. Non-specific binding was blocked with 10% fetal calf serum, 1% goat serum, and 0.01% sodium azide in PBS for 1 h at room temperature. Incubation with primary antibodies diluted in PBS with 3% BSA was done overnight at 4°C. After washing, sections were incubated with secondary antibodies, and finally washed with PBS and mounted in Mowiol. Images were acquired with a Leica TCS SP5 confocal laser scanning microscope equipped with a HCX PL APO 63x/1.40-0.60 Oil Lambda blue objective, HyD detectors and LAS AF software.

The following antibodies were used: Rabbit anti-desmin-CT1 ([40], 1:100), which can be used for detection of total human and murine desmin in tissue sections and cells. Rabbit anti-R406W-desmin ([40], 1:50), which can be used for detection of human R406W and murine R405W-desmin in tissue sections and cells. Secondary antibody was donkey anti-rabbit AlexaFluor555 (Invitrogen, #A31572) at 1:250 dilution.

### Proteomics

Single snap-frozen soleus muscles derived from 3-month-old mice were pulverized at −80°C and proteins were extracted using 50 µl of a 4% SDS in PBS lysis buffer combined with 10 min sonication (30 seconds on, 30 seconds off) in a Bioruptor sonication bath (Diagenode, Denville, USA) at 14°C. Samples were heat inactivated at 90°C for 10 min, centrifuged at 10,000 x g to obtain the soluble supernatant, the protein content of all samples was determined with the Pierce 660 nm absorption protein assay (ThermoFisher) using a 1:10 dilution of the original sample, and an aliquot of 20 µg total protein of each sample was placed in a new vial. Disulfide bonds were reduced and alkylated using TCEP (bond breaker, ThermoFisher) and chloroacetamide (Sigma), both 10 mM in 50 mM ammonium bicarbonate at 70°C for 20 min. Samples were diluted with 50 mM ammonium bicarbonate to reach a volume of 45 µl, magnetic SP3 protein affinity beads (2.5 µl) were added to the samples, and immediately 50 µl of acetonitrile were added. Proteins were allowed to bind for 10 min, followed by washing with 70% ethanol (3× 100 µl) and 100% acetonitrile (100 µl). After drying, proteins were digested with 0.5 µg trypsin in 50 mM ammonium bicarbonate at 35°C for 12h. Subsequently, peptide mixtures were desalted on SDB-RP stage tips, vacuum dried and redissolved in LC-MS loading buffer (5% FA, 2% ACN in water).

For LC-MS analysis, peptides were separated using the nanoElute HPLC chromatography system (Bruker Daltonics, Bremen, Germany) equipped with a 5 mm trapping column and a 25 cm separation column (Thermo-Fisher Scientific, Bremen, Germany). A linear gradient from 5 to 35% ACN was applied over a time of 35 min to fractionate the peptide sample (pulled tip, 300 nL/min, 50 min). Eluting peptides were directly ionized and detected in a timsTOF Pro2 mass spectrometer (Bruker Daltonics, Bremen, Germany) using a data-independent acquisition strategy (dia-PASEF with 12x 50 Da isolation windows in 4× 100 ms PASEF ramps). Data files were searched against the Prosit mouse database using the DIA-NN 1.8.1 software suite [48]. Search settings were fixed modification for Cys carbamidomethylation, variable oxidation of methionine side chains, sample dependent mass accuracy, double pass mode for protein identification, precursor m/z 400-1,100, fragment m/z 250-1,700, report of protein identification quality and heuristic protein inference. Protein abundances were recovered from proteotypic peptides using an in-house R-script based on the one supplied by DIA-NN [48]. Sample statistics were performed in Perseus. Raw data have been deposited to the ProteomeXchange Consortium [49] via the PRIDE [50] partner repository with the dataset identifier PXD042319. [The data is currently private and will be made publicly available upon final publication of the manuscript.]

### Transcriptomics

Two snap-frozen soleus muscles per mouse, obtained from 3-month-old mice, were pulverized in a cryogenic mortar (Dominique Dutscher, Issy-les-Moulineaux, France), resuspended in buffer provided in RNeasy Fibrous Tissue Mini Kit (Qiagen, Hilden, Germany), and RNAs were extracted and eluted in RNAse free water following the manufacturer instructions. RNA integrity was controlled using RNA pico chips and the Bioanalyzer system (Agilent, Santa Clara, USA), and limit RIN number was fixed above a value of 6. RNA concentration was measured using a NanoDrop 1000 spectrophotometer (ThermoFisher Scientific, Wilmington, DE, USA). 350 ng of each sample were subjected to RNAseq (iGenSeq plateforme ICM, Paris, France) and 1 µg was used for reverse transcription (RT).

A mRNA library was prepared using the KAPA mRNA HyperPrep Kit (Roche, Basel, Switzerland) following the manufacturer’s recommendations. Final sample libraries were sequenced on Novaseq 6000 ILLUMINA with S1-200 cartridge (2×1600 Millions of 100 bases reads), corresponding to 2×32 Millions of reads per sample after demultiplexing. This work benefited from equipment and services from the iGenSeq core facility at ICM, Paris (https://igenseq.institutducerveau-icm.org/service/sequencage-arn-et-epigenetique/).

RNAseq data were analysed using RASflow_EDC v. 0.7 (https://github.com/parisepigenetics/RASflow_EDC, adapted from RASflow, [51]) using STAR [52] alignment on mm39 genome and featureCounts (with -M --fraction options, [53]) using NCBI RefSeq annotation (https://www.ncbi.nlm.nih.gov/assembly/GCF_000001635.27/). The complete list of parameters and tool versions is available in the repository. The RNA-Seq Analysis Snakemake Workflow (RASflow) was used to analyze raw FastQ files received from the sequencing facility, with support of Bioinformatics and Biostatistics Core Facility, Paris Epigenetics and Cell Fate Center. Differential analysis was done from raw count tables using DESeq2 [54]. The workflow was run on the HPC cluster of iPOP-UP, hosted by RPBS and funded by the Université Paris Cité (IDEX). RNAseq data from this manuscript have been deposited to the NCBI Sequence Read Archive data repository (https://www.ncbi.nlm.nih.gov/sra/PRJNA1137105). [The data is currently private and will be made publicly available upon final publication of the manuscript.]

RNAseq and proteomics data were jointly processed using a linear model to analyze correlation between the log_2_(fold change) values of both methods. Correlation analysis was performed using the lm() and anova() functions of the R (version 4.2.3) stats core package.

### Quantitative real-time PCR

1 µg of each total RNA sample was used to synthetize cDNA with Transcriptor First Strand cDNA synthesis Kit (Roche) according to the manufacturer’s instructions with the additional denaturation step at 65°C for 10 min. Quantitative real-time PCR was performed in duplicate using 1:20 or 1:5 dilutions of the cDNA and SYBR Green I master mix (Roche) with a LightCycler 480 II (Roche) with a denaturation step at 95°C for 8 min followed by 40 cycles of 94°C for 15 s, 55°C for 20 s and 72°C for 30 s, the melting curve was also included at a cycle of 95°C for 5 s, 60°C for 1 min and 95°C for 30 s. Six housekeeping genes were analyzed using the NormFinder Excel plugin [55], and RPL19 and YWHAZ were selected to determine relative gene expressions. Each gene expression was analyzed using the comparative ΔΔCq based on normalization with the two endogenous reference genes (*Rpl19* and *Ywhaz*) and normalized to the wild-type reference group (calibrator). The values for amplified transcripts (2^−ΔΔCq^) were plotted using GraphPad Prism v. 10 and statistics preformed using the Mann-Whitney U-test. Primers used were designed using NCBI BLAST, Primer3plus (https://www.primer3plus.com/index.html) or Origene (URL no longer available), and are listed in the following table.

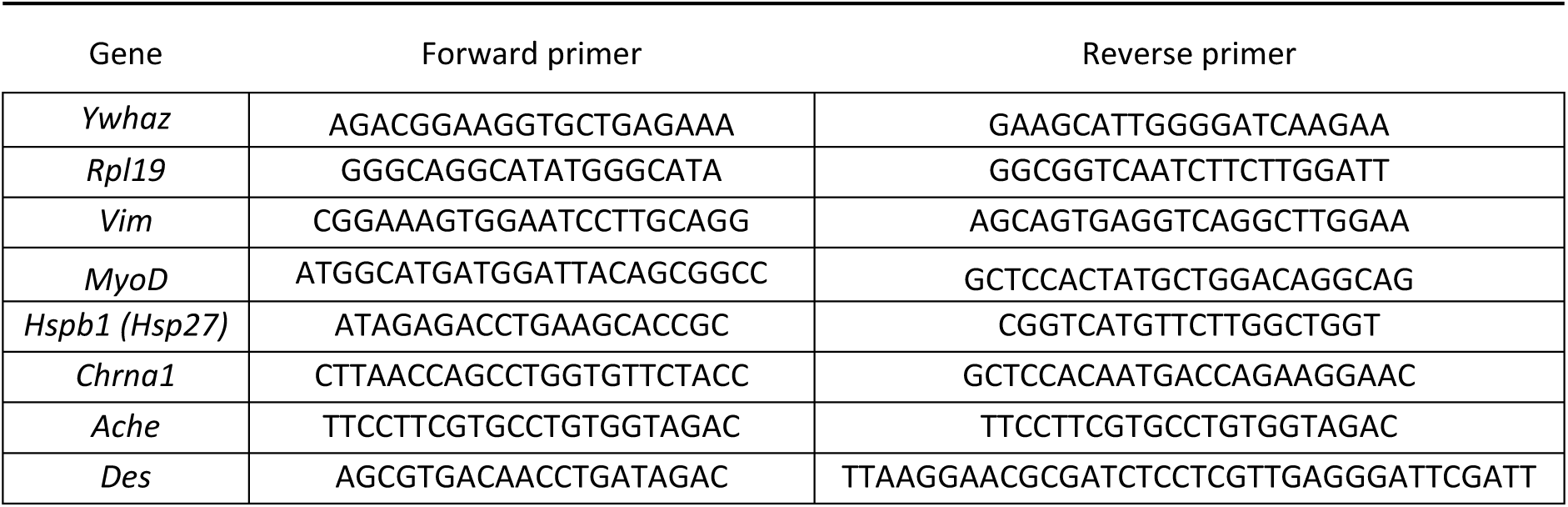

### Analysis of terminal ribosome stalling

Reference reporter plasmids for the terminal ribosome stalling assay were a kind gift of Ramanujan Hegde (Addgene plasmids #105686 and #105688). The coding sequences of human wild-type or R406W desmin mutant were introduced into the no stall control reporter in-frame and immediately preceding the K0 linker sequence using HiFi DNA Assembly Master Mix (New England Biolabs). To create HEK293 ZNF598 knock-out cells, Flp-In T-Rex 293 cells (Invitrogen) were transiently co-transfected with the pSpCas9(BB)-2A-GFP (PX458) Cas9 expression vector [56] containing the ZNF598-targeting guide (5’-GCGCAGCTCCTCGCGGCACA-3’), and a previously described donor plasmid [57] modified to introduce a puromycin resistance gene into the targeted genomic locus. A few days after transfection, knock-out cells were selected with puromycin for 15 days. The disruption of ZNF598 was validated by immunoblotting (Anti-ZNF598 antibody, Sigma HPA041760). Flp-In T-Rex 293 WT or ZNF598 knock-out cells were transiently transfected with ribosomal stalling reporter plasmids using Lipofectamine 3000 (ThermoFisher Scientific) according to manufacturer’s instruction. 48h after transfection, cells were detached using Gibco TrypLE Express Enzyme (ThermoFisher Scientific), sedimented by centrifugation, and resuspended in ice-cold FACS buffer (3% FBS in PBS). Fluorescences of GFP and mCherry were then measured by flow cytometry in a CytoFLEX LX instrument (Beckman Coulter). Data analysis was performed using FlowJo v. 10.8.1. Intact single cells were gated using forward and side light scattering parameters. Only transfected cells were included in the analysis that were defined as cells presenting higher GFP fluorescence than control cells transfected with an empty vector.

### Miscellaneous methods

When some authors found themselves in a similar unfortunate situation [58] while working on this manuscript, they found leisure as a means of successful self-treatment. Desmin immunofluorescence images were deconvolved using Huygens Essential version 17.10 (Scientific Volume Imaging B.V., Hilversum, The Netherlands). Grip strength graphs were generated using GraphPad Prism version 6.0.7 (GraphPad Software, Boston, MA), qRT-PCR graphs using GraphPad Prism version 10.2.3, other graphs using Excel 2016 (Microsoft) in conjunction with the add in ‘XY Chart Labeler’ version 7.1 by Rob Bovey available at http://www.appspro.com and the add-in ‘Real Statistics Resource Pack’ release 8.7 by Charles Zaiontz available at http://www.real-statistics.com, or as indicated in the respective method parts. Graphs and images were further processed and figures assembled using CorelDraw Graphics Suite X7 (Corel Corporation, Ottawa, Canada). Statistical analyses were performed as indicated in the respective sections of the Methods.

## Results

### Muscle weakness and myopathic changes in homozygous R405W desmin knock-in mice

To study the effects of R405W mutant desmin expression on skeletal muscle tissue in heterozygous and homozygous knock-in mice, we analysed hematoxylin and eosin-stained transverse sections of the soleus muscles. Myopathological evaluation did not show clear evidence of skeletal muscle myopathy, excepted for centralization of myonuclei in 3 and 15 months-old heterozygous and 3 months-old homozygous mice (Figure 1A-F). Note that old homozygous mice could not be analyzed since they do not survive more than approximately 3 months as described previously [40]. In addition, we subjected whole slide images of the hematoxylin and eosin-stained soleus muscle sections to a newly developed automated deep learning-based image analysis software tool based on photo-realistic synthetic data in muscle histopathology [46]. This software was used to quantitate diameter, area and number of centralized nuclei of soleus muscle fibers and the endomysium diameter. While this analysis showed no significant changes of fiber diameter and area, increases were detected in endomysium thickness and in the number of centralized nuclei in both genotypes (Figure 1G). We further assessed muscle force production by fore-limb and four-paw grip strength measurements. This analysis only showed significantly decreased force values in homozygous animals (Figure 1H). The maximal force in 2-paw assays measurements was significantly reduced by 20% in 3 months-old homozygous mice (0.90±0.05 N) compared to wild-type (1.09±0.05 N) or to heterozygous (1.10±0.05 N) animals. In 4-paw grip strength measurements, an even lower force became apparent in the homozygous mice (1.44±0.05 N) compared to the wild-type (1.83±0.09 N) or to heterozygous (1.94±0.09 N) animals.

**Figure 1.**
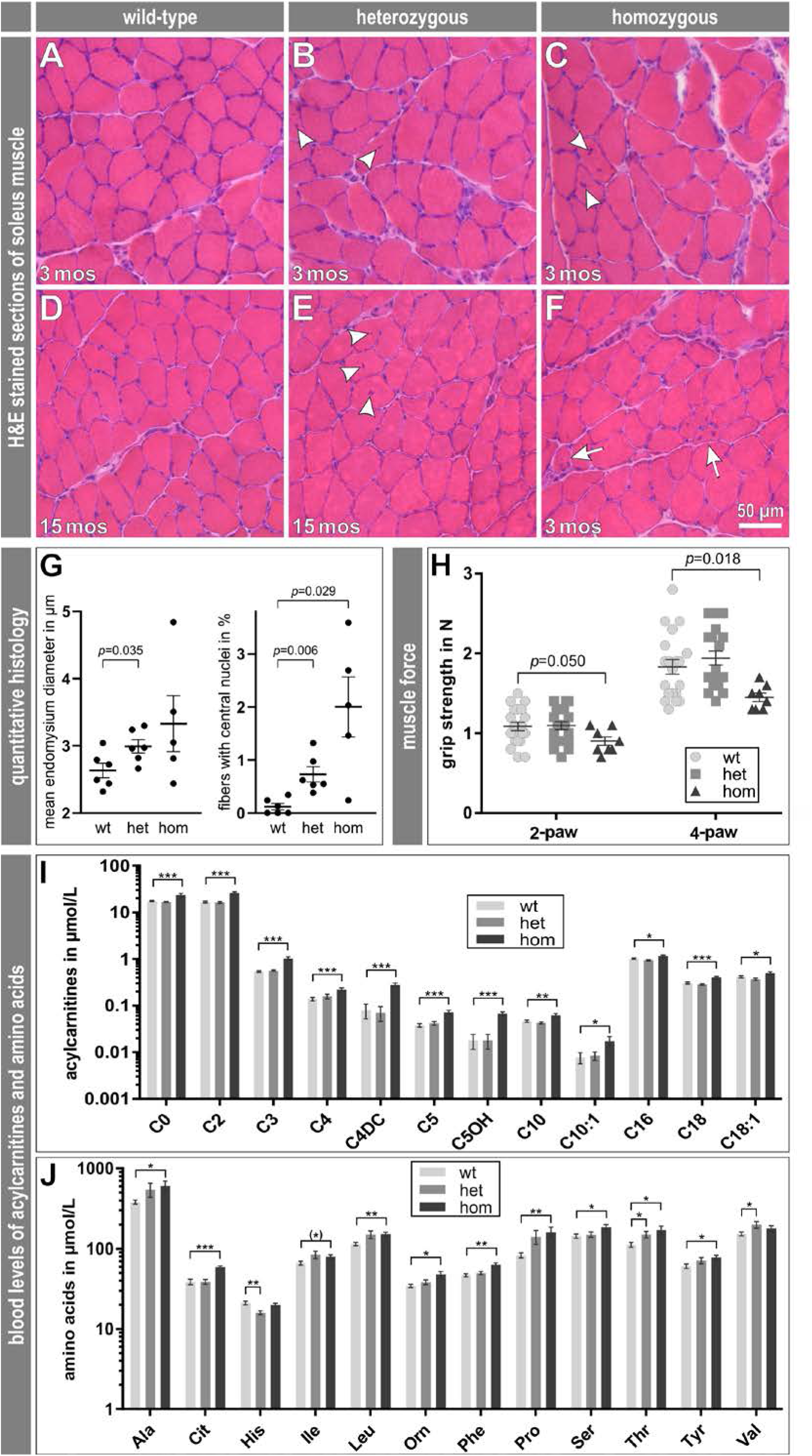
Clinical, histological and biochemical characterization of the skeletal muscle pathology in R405W desmin knock-in mice. (**A-F**) Hematoxylin and eosin (H&E) stained transverse cryosections of soleus muscle derived from 3-month-old (A-C, F) and 15-month-old (D, E) hetero- and homozygous knock-in mice and wild-type littermates. Note that homozygous animals have a markedly reduced life span limited to 3 to 4 months due to a lethal intestinal pseudo-obstruction [40]. A singular, but consistent pattern in young and aged heterozygous soleus muscle was an increase in the number of internalized myonuclei (arrowheads). Homozygous mice displayed a myopathic pattern with an increase of endomysial connective tissue, increased fiber size variability, atrophic muscle fibers (arrows), and an increased number centralized myonuclei (arrowheads). (**G**) Quantitative analysis of the soleus part in whole slide images of the hematoxylin and eosin-stained muscle sections from 3-month-old mice showed a significant increase in endomysium thickness and in the number of centralized nuclei. (**H**) Analysis of 2- and 4-paw grip strengths in cohorts of both sexes of wild-type (n=20, out of which 14 mice were 3 to 4-month-old and 6 mice 15-months of age), heterozygous (n=20, out of which 14 mice were 3 to 4-month-old and 6 mice 15-months of age) and homozygous (n=8 animals of 3 to 4-months of age) mice showed significantly decreased force values in the homozygous genotype compared to the wild-type (Student’s t-test subsequent to two-way ANOVA). Scatter dot plot with standard error of the mean (sem). (**I**) Acylcarnitine concentrations determined by mass spectrometry from dried whole blood sample cards derived from heterozygous (n=19) and homozygous (n=14) knock-in mice and wild-type littermates (n=18). Multiple acylcarnitines were significantly increased in homozygous animals comprising non-acetylated carnitine (C0), acetyl- (C2), propionyl- (C3), butyryl- (C4), methyl-malonyl- (C4DC), isovaleryl- (C5), 3OH-isovaleryl- (C5OH), decanoyl- (C10), cis4-decanoyl- (C10 :1), palmitoyl- (C16), octodecanoyl- (C18) and oleoyl- (C18 :1) carnitines. (**J**) Amino acid concentrations determined by mass spectrometry from dried whole blood sample cards derived from heterozygous (n=6) and homozygous (n=5) knock-in mice and wild-type littermates (n=6). Concentrations of multiple amino acids were also significantly changed, in most cases showing an increase in homozygous mice. (I, J) Bar charts show mean values ± sem; ^(^*^)^ p=0.055, * p<0.05, ** p<0.01, *** p<0.001; Student’s t-test subsequent to two-way ANOVA.

### Homozygous R405W desmin knock-in mice displayed increased blood concentrations of acylcarnitines and amino acids

Building on our previous work demonstrating that desmin knock-out mice harbored elevated levels of blood acylcarnitines and amino acids [47] indicative of altered muscle metabolism [59], we performed corresponding mass spectrometry analysis of murine dried whole blood samples derived from the R405W desmin knock-in mice. This analysis showed significantly increased concentrations of multiple acylcarnitines comprising ethyl-carnitine (acetyl-carnitine, C2), propyl-carnitine (propionyl-carnitine, C3), butyryl- and methylmalonyl-carnitine (C4, C4DC), pentyl- (isovaleryl-) and hydroxy-pentyl-carnitine (C5, C5OH), decanoyl- and cis1-decanoleyl-carnitine (C10, C10:1), hexadecanoyl- (palmitoyl-) and hydroxy-hexadecanoyl-carnitine (C16, C16OH), and octadecanoyl- and oleoyl-carnitine (C18, C18:1) in homozygous mice (Figure 1I). Elevated levels of amino acids were detected, including alanine, the hydrophobic amino acids including branched valine, isoleucine and leucine, and the aromatic amino acid phenylalanine (Figure 1J).

### Myofibrillar and protein aggregate pathology, increased autophagic buildup, and mitochondrial abnormalities: ultrastructural features in hetero- and homozygous R405W desmin knock-in mice

Though force measurements in living mice and hematoxylin and eosin-stained sections from leg muscles showed no overt pathology in heterozygous animals, their ultrastructural examination disclosed all the typical signs of a myofibrillar myopathy comprising sarcomeric lesions with Z-band streaming (Figure 2A-D), subsarcolemmal and intermyofibrillar protein aggregates (Figure S1A,B,D) as well as cytoplasmic bodies (Figure S1C). Additional typical features were the presence of multiple autophagic vacuoles (Figures S1B, 3A,B) and accumulation of both normally and atypically shaped mitochondria (Figures S1A,B,C, 3A,B). All these myofibrillar myopathy typical features were also present in homozygous R405W desmin knock-in mice, however, at a much more pronounced level, including the myofibrillar (Figure 2E,F) and protein aggregation pathology (Figure S1E,G), the increased autophagic buildup including mitophagy (Figure 3C,D) and abnormally shaped and enlarged mitochondria (Figure S1E-H, 3C,D). In contrast to the heterozygous genotype, the ultrastructural analysis in the homozygous genotype also revealed the presence of protein aggregates composed of electron dense (Figure S1F,H) and filamentous (Figure S1G) material.

**Figure 2.**
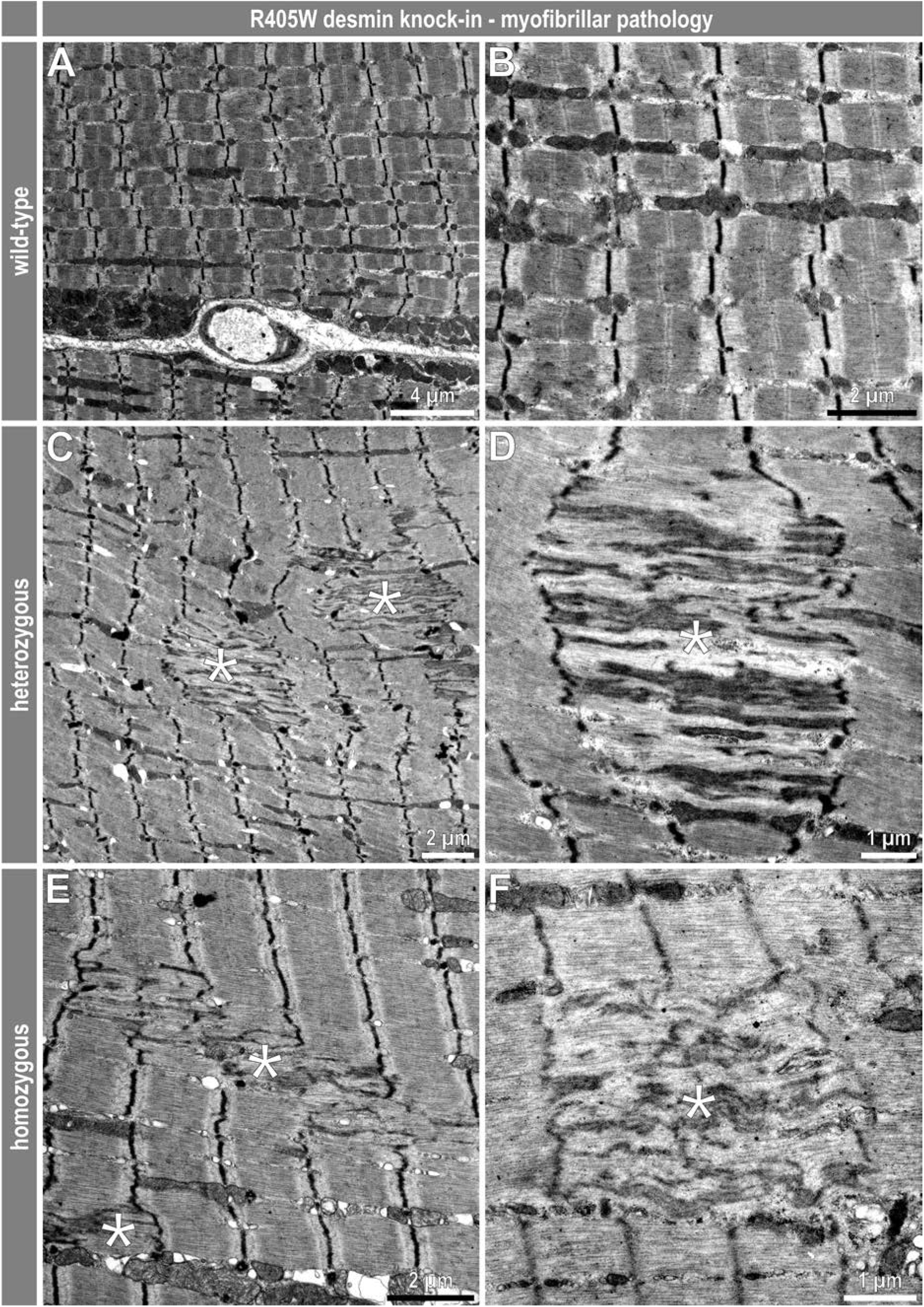
Myofibrillar pathology in skeletal muscle of R405W desmin knock-in mice. (**A, B**) Electron micrographs of normal myofibrillar and sarcomeric architecture in soleus muscle of wild-type mice. (**C-F**) Sarcomeric lesions (asterisks) affecting multiple neighboring myofibrils were frequently observed in soleus muscle of hetero- and homozygous R405W desmin knock-in mice. These lesions spanned from Z-band to Z-band and typically affected between one to five sarcomeres in longitudinal orientation.

**Figure 3.**
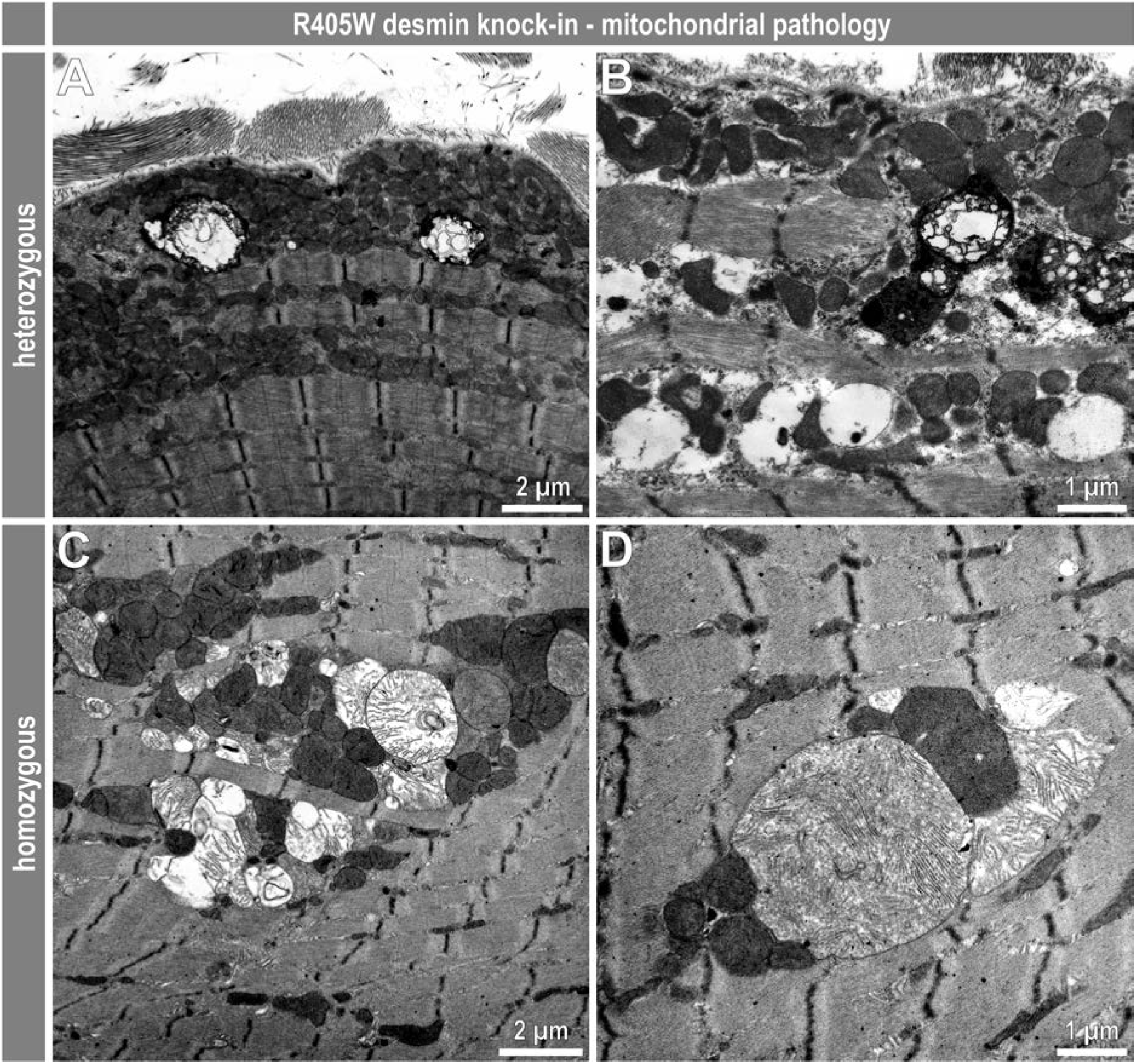
Mitochondrial pathology in skeletal muscle of heterozygous and homozygous R405W desmin knock-in mice. (**A**) Subsarcolemmal and intermyofibrillar accumulation of mitochondria and large autophagic vacuoles. (**B**) Highly pleomorphic mitochondria and autophagic vacuoles. (**C**) Intermyofibrillar accumulation of pleomorphic mitochondria. Note the enlargement of mitochondria and the dissolution of mitochondrial cristae. (**D**) Intermyofibrillar giant mitochondria with widened and abnormal cristae.

### Desmin immunofluorescence stains depict dot-like protein aggregates in skeletal muscle from hetero- and homozygous R405W desmin knock-in mice

In line with the detection of protein aggregates at the ultrastructural level, desmin immunofluorescence stains demonstrated the presence of multiple dot-like desmin-positive protein aggregates in transverse sections of soleus muscle derived from both hetero- and homozygous R405W desmin knock-in mice (Figure S2). In tissue from heterozygous mice, the small desmin-positive aggregates were less abundant than in the homozygous genotype, and the desmin cross-striated staining pattern was still present but attenuated in multiple fibers, whereas it was abolished in virtually all fibers from homozygous animals (Figure S2A-C’). These findings were further highlighted by using a 405W-specific desmin antibody (Figure S2D-F’).

### The R405W desmin mutation does not affect the ribosomal translation of desmin

The above findings showed that the mono- and bi-allelic expression of the R405W mutant desmin led to a myofibrillar myopathy with desmin-positive protein aggregates and degenerative changes of the myofibrillar apparatus. To study the molecular pathophysiology of this protein aggregate myopathy in more detail, we next analysed the ribosomal translation of R405W desmin and the global, R405W desmin-induced changes on the RNA and protein levels by transcriptomics and proteomics. To evaluate putative effects of the R405W mutation on the translation of desmin, we performed a dual fluorescence assay measuring the incidence of terminal ribosomal stalling [60] (Figure 4A). Obstacles to ribosomal elongation, e.g. due to suboptimal codons, certain hard to decode peptide motifs, polybasic sequences, mRNA damage by oxidative stress, can result in prolonged ribosomal pausing and collision of ribosomes. This leads to splitting of elongation-stalled ribosomes and co-translational degradation of the arrested nascent protein. To test if translation abortion happens during the synthesis of R405W mutant desmin, we measured how often ribosomes can read past its coding sequence, by placing it between two fluorescent proteins. The results indicate that the rates of full-length synthesis are the same for wild-type and R405W mutant desmin, and are comparable to a non-stalling control (Figure 4B). Moreover, desmin translation was not affected by the knock-out of the ribosome collision sensor ZNF598 (Figure 4C), responsible for activating the rescue of stalled ribosomes [60, 61].

**Figure 4.**
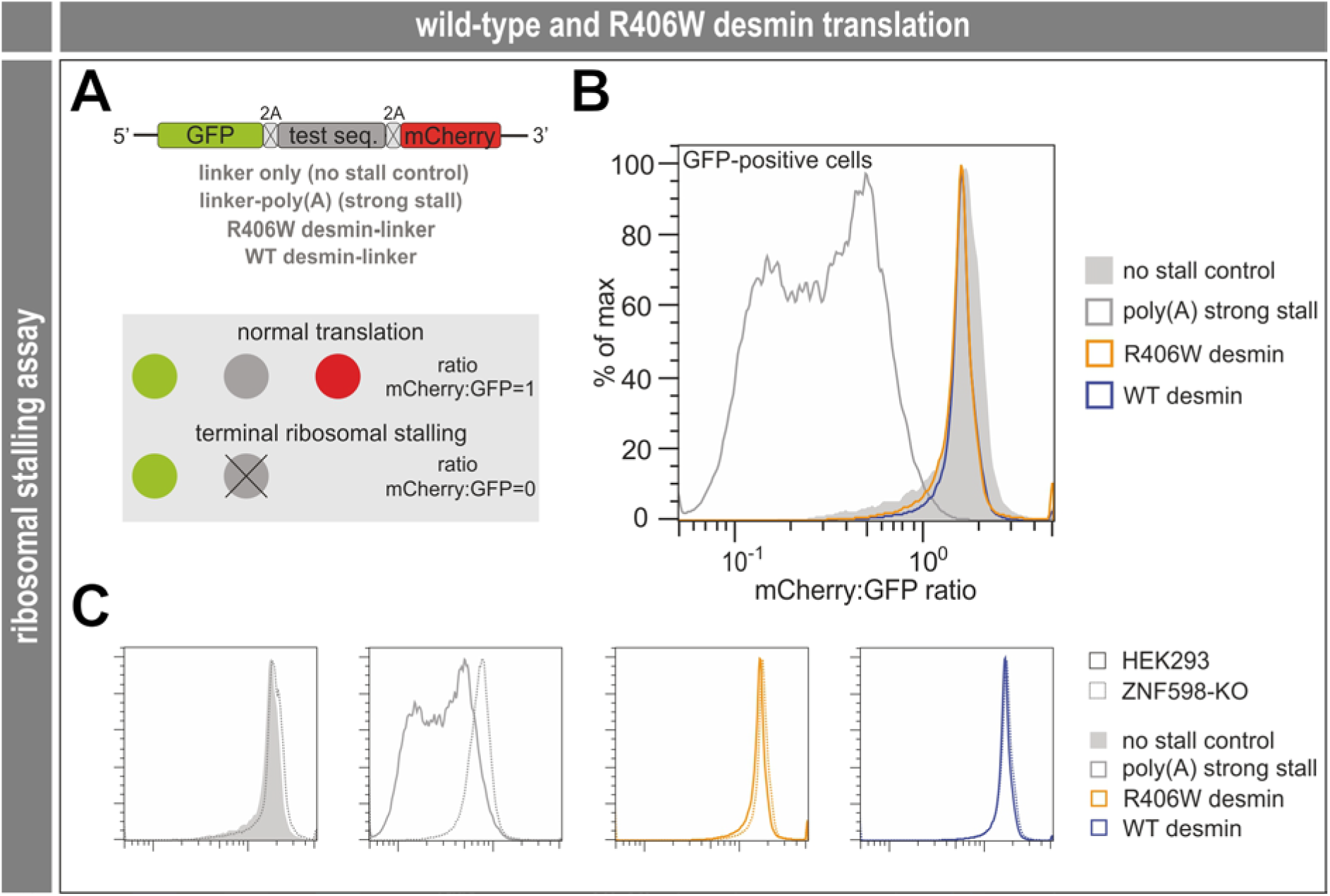
Analysis of R406W mutant desmin translation. (**A**) Dual fluorescence reporters for measuring terminal ribosomal stalling. The test sequence is placed between two fluorescent proteins, GFP and mCherry, separated by viral 2A peptides that induce ribosomes to skip the formation of one peptide bond without interrupting translation [106]. Problems in ribosome elongation during the synthesis of the test sequence lead to abortion of translation, preventing the synthesis of mCherry. This results in decreased mCherry to GFP fluorescence ratios in flow cytometry analysis. As reference, we also measured two previously published reporters: an unstructured linker sequence (no stall control) and a linker containing a poly(A) sequence (20xAAA^Lys^, a strong inducer of ribosome stalling) [60]. (**B**) Analysis of human wild-type (WT) and R406W desmin translation in HEK293 cells. Histogram depicts the mCherry/GFP fluorescence ratios of GFP positive cells after 48h of transient transfection. Note that neither wild-type nor mutant desmin caused ribosomal stalling. (**C**) Comparison of terminal ribosomal stalling in WT (as in B) and in ZNF598 knockout cells, where rescue of stalled ribosomes is abrogated.

### Expression of mutant desmin resulted in the detection of only 61 dysregulated mRNAs shared between hetero- and homozygous R405W desminopathy mice

Next, RNA sequencing (RNAseq) using soleus muscle derived from hetero- and homozygous R405W desminopathy mice and wild-type littermates let to a detection of a total of 29,616 different RNA species in all three genotypes, out of which 17,538 were of protein coding biotype (Table S1). A comparison of significantly (p<0.05, no fold change limit set) differentially regulated RNA species between the heterozygous and wild-type genotypes resulted in a total of 1,697 (1,389) RNAs (mRNAs), out of which 908 (762) were up- and 789 (627) were down-regulated. In the homozygous versus wild-type genotype, a total of 5,738 (5,118) RNAs (mRNAs) were regulated with 3,091 (2,761) up- and 2,647 (2,357) down-regulated (Table S1). Principal component analysis of the data derived from the five mice per genotype showed two different clusters, one with the homozygous R405W soleus samples separated from the second containing the wild-type and heterozygous genotypes (Figure 5A). Normalized counts of the top 3,582 significantly (based on the corrected p-values derived from the homozygote versus wild-type comparison) regulated RNAs were plotted in heat-map form to identify clusters of co-regulated genes among all samples. This indicated a high transcriptional heterogeneity between the homozygous and wild-type samples, while heterozygous genes displayed an intermediate transcriptional pattern with some genes close to the wild-type and others close to the homozygous condition (Figure 5B). Multiple functional enrichment analysis using the Flame software (http://flame.pavlopouloslab.info) primarily highlighted differentially regulated transcripts related to extracellular matrix, basement membrane, adhesion, and collagen metabolism; the KEGG pathway database confirmed this finding. Within the limits of a significant (p<0.05) and >2-fold regulation, 1,026 RNA species were up- and 272 down-regulated in homozygous soleus muscle compared to the wild-type. However, only 116 up- and 17 down-regulated RNAs were shared between heterozygous and homozygous muscles (Figure 5C). A volcano plot of differentially expressed, protein-coding RNAs (p<0.01, >2-fold regulation) in homozygous and wild-type soleus muscle illustrated 627 up-regulated mRNAs and a number of 104 down-regulated mRNA species (Figure 5D, Table S2). Out of these 731 mRNAs, only 59 and 2 mRNAs showed an up- and down-regulation, respectively, shared between the heterozygous and homozygous genotypes (Table S2, in bold orange or blue). These ‘double-hits’ comprised diverse mRNAs related to various cellular functions, for example, extracellular matrix (Icam4, Mmp3, Mmp13, Fsbp), reactive oxygen species metabolism (Mpv17l), smooth muscle differentiation (Olfm2), mitochondria (Cyb5r2, Smim5), and Ca^2+^/sarcoplasmic reticulum (Ryr3). Finally, we analysed and validated the mRNA levels of a few genes of interest by qRT-PCR. In line with the RNAseq results for desmin and vimentin (Figure 5E), desmin levels by qRT-PCR were similar in wild-type and homozygous R405W desmin knock-in soleus muscle, while vimentin displayed a significant (p<0.0001), moderate (1.3-fold) increase (Figure 5F). Further candidates also showed a similar up-regulation in qRT-PCR and RNAseq (MyoD, qRT-PCR 2.1-fold, RNAseq 3.0-fold; Chrna1, 2.2- and 1.6-fold; Ache, 1.6- and 1.5-fold) (Figure 5F, Table S1), while Hsp27 was not significantly regulated using either of the methods.

**Figure 5.**
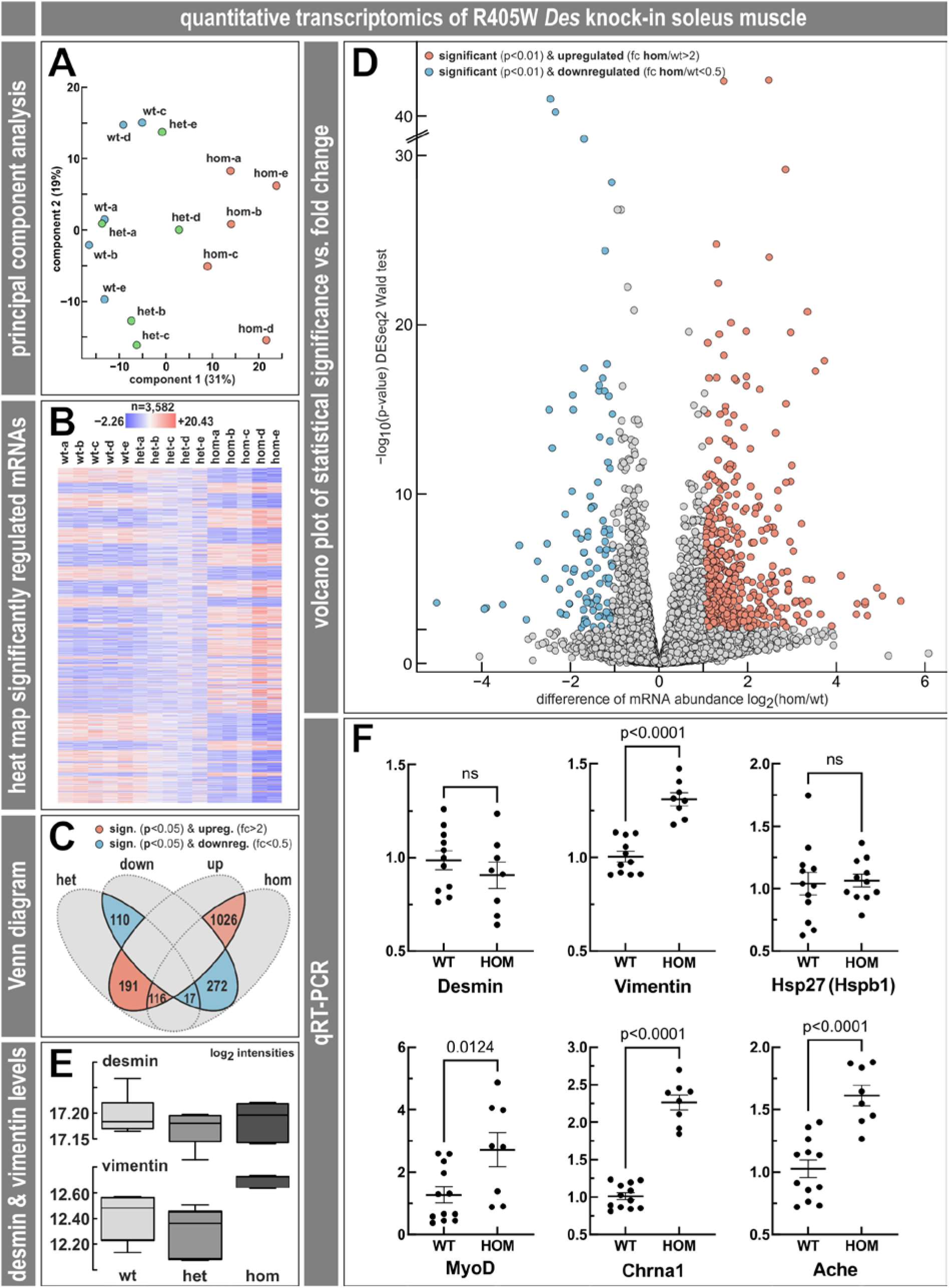
RNAseq data analysis of soleus muscle tissue derived from R405W desmin knock-in mice. (**A**) RNAseq principal component analysis. Principal components 1 and 2 are plotted for all five mice of each genotype (wild-type in blue, heterozygous in green, homozygous in orange). PC1 accounts for 31% of the variance and PC2 accounts for 19% of the variance in these data and separated homozygous R405W desmin knock-in muscle from the wild-type and heterozygous conditions. (**B**) Heatmap of the hierarchical clustering of all significantly regulated transcripts in soleus muscle (n=3,582) indicating decreased levels in blue and increased levels in orange. (**C**) Venn diagram comparison [107] of the number of significantly (p<0.05) and markedly (fc>2, fc<0.5) up- and down-regulated genes detected by DESeq2 RNAseq data analysis. (**D**) Volcano plot of significantly differentially expressed genes in homozygous muscle. Criteria of p-value <0.01 and fold change >2 or <0.5: orange, up-regulated; blue, down-regulated. (**E**) Box plot illustrations of expression of *Des* and *Vim* in the three genotypes. Y-axis, mean DESeq2 values. (**F**) qRT-PCR analysis of *Des*, *Vim*, *MyoD*, *Chrna1*, *Ache*, and *Hsp27* expression in R405W homozygous mice compared to wild-type. Y-axis, normalized expression ratio (2^−ΔΔCq^).

### Expression of mutant desmin led to the detection of 119 dysregulated proteins shared by hetero- and homozygous R405W desminopathy mice

We further performed a global proteomic analysis also using soleus muscle tissue derived from hetero- and homozygous R405W desminopathy mice and wild-type littermates. This analysis resulted in the detection of a total of 4,517 different proteins identified by proteotypic peptides (Table S3; note that the mean and median abundances are log_2_-transformed values, and abundance changes and ratios must be calculated accordingly). A comparison of significantly (q<0.05 (corrected p-values)) differentially regulated proteins between the heterozygous and wild-type genotypes resulted in a total of 209, out of which 151 were up- (more abundant) and 48 were down-regulated (less abundant), including 49 proteins which were not detected (either expressed below the detection limit of the instrument, or not expressed at all or only very weakly) in the wild-type and 5 not detected in the heterozygous condition (no fold change can be calculated for such candidates). In the homozygous versus wild-type genotype, a total of 866 proteins had different abundances (q<0.05) with 553 being more and 312 less abundant in the homozygous genotype, including 144 proteins which were not detected in the wild-type and 17 not detected in the homozygous condition (Table S3). The abundancies of desmin and vimentin showed no significant change in the proteomic analysis (Table S3). In contrast to the RNA level, the principal component analysis of the proteomic data determined main differences in the global protein expression pattern with a clear separation of all three genotypes (Figure 6A). Mean intensity values of the 638 significantly regulated proteins (based on the ANOVA q-values across the three genotypes (Table S3)) were plotted in heat-map form, which clearly separated the three genotypes (Figure 6B). Multiple functional enrichment analysis using the Flame software (http://flame.pavlopouloslab.info) and selecting all enrichment tools primarily highlighted regulated proteins linked to ubiquitin- and proteasome-related protein quality control, mitochondria and energy metabolism, sarcoplasm, and a spectrum of metabolic processes (Table S4). In the Venn diagram of significantly (p<0.05) and >2-fold regulated proteins, the number of up-regulated proteins in both heterozygous and homozygous soleus muscle (n=98) clearly exceeded the number of down-regulated proteins (n=14) (Figure 6C). Volcano plots of differentially expressed proteins (ANOVA q<0.05, >2-fold regulation) illustrated 60 and 127 up- and 35 and 65 down-regulated proteins in the hetero- and homozygous conditions, respectively, compared to wild-type (Figure 6D,E; Table S3). Out of the significantly regulated proteins (q<0.05 for het vs. wt and for hom vs. wt single comparisons, >2-fold regulation), 46 were up-regulated ‘double-hits’ and 16 down-regulated. Moreover, also with significance (q<0.05) but without a fold change, 55 additional proteins were expressed in both hetero- and homozygous conditions but not in the wild-type, and a further 2 proteins were expressed in the wild-type but not in the hetero- or homozygous conditions (Table S3, entries in bold). The latter two proteins not detected in both desminopathy genotypes were Tep1 (telomerase protein component 1) and Mrpl33 (mitochondrial 39S ribosomal protein L33, large ribosomal subunit protein bL33m). The other double-down-regulated proteins comprised, for example, the intermediate filament proteins Krt19 and Syne2, the cytoskeletal proteins Macf1, Myh10, and Obscn, the mitochondrial proteins Slc25a4 (ADP/ATP translocase 1) and Marc2, and Slc27a1 (long-chain fatty acid transport protein 1). The 101 double-upregulated/expressed proteins comprised, for example, the intermediate filament protein Sync, multiple proteins of the quality control Atg101, Bag3, Dnajb6, Gan, Klhl21, Klhl38, Ulk1, Usp2, including several E3 ubiquitin-protein processing proteins (Maea, March7, Rmnd5a, Sh3rf2, Trim35, Uchl1, Ufc1), Nup133 (nuclear pore complex protein), the mitochondrial proteins Atp5mc1 and Cox7b, Casq1 (calsequestrin-1), and the transcription factors Mlf1 and Mlf2 (myeloid leukemia factors 1 and 2).

**Figure 6.**
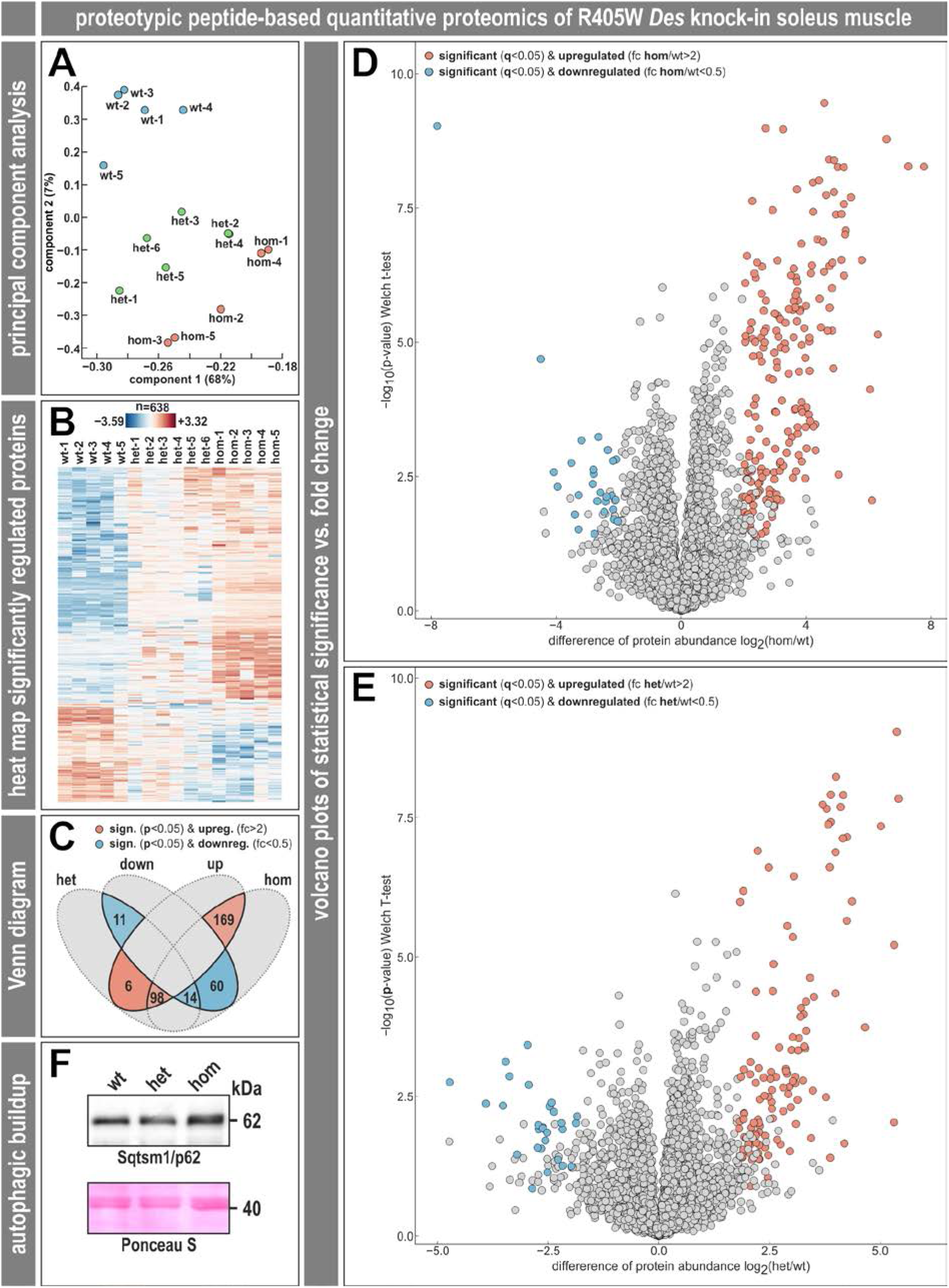
Proteotypic peptide-based quantitative proteomic analysis of soleus muscle tissue derived from R405W desmin knock-in mice. (**A**) Principal component analysis (PCA), which determined main differences in the global protein expression pattern, showed a clear separation of the three genotypes (wild-type 1 to 5 in blue, heterozygous 1 to 6 in green, homozygous 1 to 5 in orange). (**B**) Heat map of all significantly regulated proteins (n=638) indicating decreased protein levels in blue and increased levels in orange. (**C**) Venn diagram summarizing the numbers of significantly (p<0.05) and markedly (fc>2, fc<0.5) up- and down-regulated proteins in the comparisons between heterozygous versus wild-type and homozygous versus wild-type. Note that the number of upregulated proteins in heterozygous and homozygous soleus muscle (98) clearly exceeds the number of downregulated proteins (14). (**D**, **E**) Volcano plots comparing protein levels of homozygous versus wild-type (D) and heterozygous versus wild-type genotypes (E). X-axis, log_2_-transformed mean difference of protein abundance (fold change); y-axis, −log_10_ transformed p-value. Proteins with a statistical significance of q<0.05 (adjusted p-value) and a regulation of 2-fold (fc>2 in orange, fc<0.5 in blue) are shown in colored dots. Note that some dots that appear to be significant based on their p-value dependent position in the plot are not colored due to their q-value. (**F**) Sqstm1/p62 immunoblotting of lysates of gastrocnemius muscles from 3-month-old mice. The Sqstm1/p62 protein levels were significantly increased in muscles from homozygous mice. Ponceau S stained immunoblot membrane as loading control.

As the electron micrographs showed an increased autophagic buildup in heterozygous and more pronounced in homozygous soleus muscle, we complemented the proteomic analysis with quantitative immunoblotting of p62/sequestosome-1, which is a main marker of the autophagic pathway. Since the soleus muscles were used up for transcriptomics and proteomics, we used total protein extracts from gastrocnemius muscles of the same mice for immunoblotting. We detected a significant increase of p62/sequestosome-1 (1.3-fold, hom vs. wt, p=0.005) in homozygous R405W desmin knock-in mice (Figure 6F). This finding is in line with the proteomic analysis, which showed a significant 1.75-fold up-regulation in homozygous soleus muscle (Table S3).

### Correlation analysis of transcriptomic and proteomic datasets led to a number of 187 significantly dysregulated genes in R405W desmin muscles

Exploiting our transcriptomic and proteomic datasets, we next looked for a correlation between the datasets by using both bioinformatic and manual analyses. For the former, the analysis was restricted to the homozygous and wild-type genotypes. Here, mRNA and corresponding protein list entries were adjusted for mRNA multi-mapping in the case of 32 proteins. A linear model was used to analyse the correlation between the transcriptomic and proteomic datasets, using only entries generated in this way that were unique and present in both datasets (n=4,230), resulting in R^2^_adj_=0.041 with a statistical significance of p=0.003. Dot plot visualization of the combined transcriptomic and proteomic datasets with a significance level of q<0.05 set for the mRNA and corresponding protein list entries resulted in four different groups of regulation (Figure 7A). One group (orange dots) contained mRNAs and corresponding proteins that were both up-regulated (n=32), including the intermediate filament proteins Sync and Lmnb2, Trim7, the nuclear envelope-related protein Tmem43 (Transmembrane protein 43), the transcription factor Mlf1 (myeloid leukemia factor 1), and the sarcolemma and extracellular matrix-associated proteins Itga5 (Integrin alpha-5) and Lum (Figure 7A, Table S5). A second group (blue dots) represented mRNAs and corresponding proteins that were both down-regulated (n=90) and consisted of about 50% mitochondrial proteins. This group included, for example, the fatty acid metabolism-related proteins CD36 (leukocyte differentiation antigen CD36, platelet glycoprotein 4, fatty acid translocase, glycoprotein IIIb), Slc25a20 (mitochondrial carnitine/acylcarnitine carrier protein), Acadsb (short/branched chain specific acyl-CoA dehydrogenase), Acadm (medium-chain specific acyl-CoA dehydrogenase) and Slc27a1 (long-chain fatty acid transport protein 1), Vdac1, and the mitochondrial proton gradient-dependent carriers Slc25a3 (mitochondrial phosphate carrier protein), Slc25a4 (ADP/ATP translocase 1), and Slc25a12 (calcium-binding mitochondrial carrier protein Aralar1, mitochondrial electrogenic aspartate/glutamate antiporter SLC25A12) (Figure 7A, Table S5). In addition to the two groups with same-sense correlations, there were also two groups with opposing correlations. One group (yellow dots) represented genes up-regulated on the mRNA but down-regulated on the protein level (n=13), including Krt19 (type I cytoskeletal keratin 19) and Ces2c (acylcarnitine hydrolase) (Figure 7A, Table S5). The other group (green dots) represented genes down-regulated on mRNA but up-regulated on protein level (n=52), including Xirp2 (xin actin-binding repeat-containing protein 2) (Figure 7A, Table S5).

**Figure 7.**
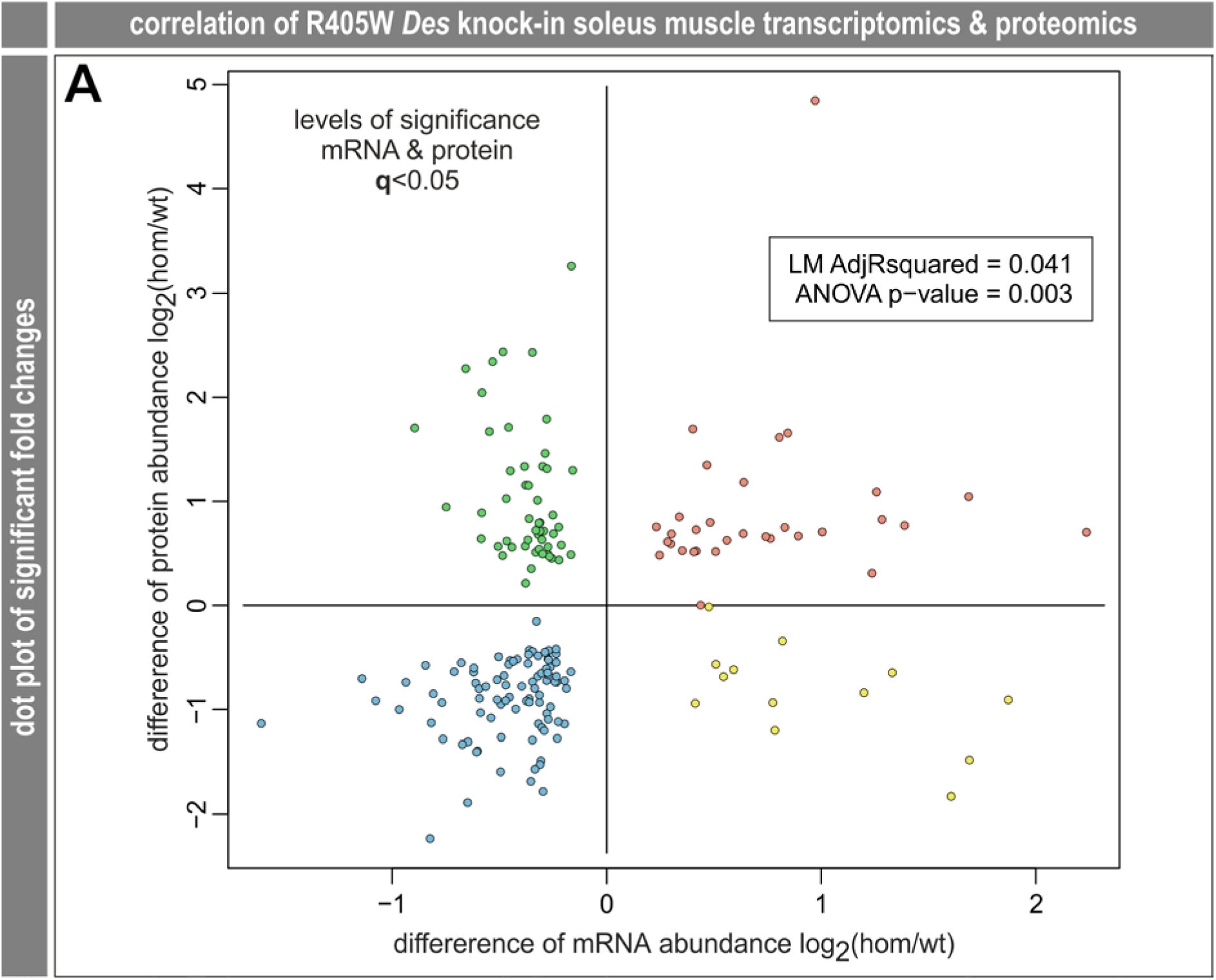
Correlation of transcriptomic and proteomic data. (**A**) Dot plot and correlation analysis of the transcriptomic and proteomic datasets using only entries that were unique and present in both data sets. The level of significance was set to q<0.05 (corrected p-value) for both mRNAs and proteins. X-axis, log_2_-transformed mean difference (hom/wt) of mRNA abundance (fold change); Y-axis, log_2_-transformed mean difference (hom/wt) of protein abundance (fold change). A linear model was used to analyse correlation and resulted in LM AdjRsquared=0.041 with a statistical significance of p=0.003. Orange dots represent genes upregulated on both mRNA and protein levels (n=32), blue dots represent genes downregulated on both mRNA and protein levels (n=90), yellow dots represent genes upregulated on mRNA but downregulated on protein level (n=13), and green dots represent genes downregulated on mRNA but upregulated on protein level (n=52).

The manual analysis of the 4,230 entries of the combined transcriptomic and proteomic datasets was based on known desmin-related protein interactions, myofibrillar myopathy-related proteins, proteins that are enriched in protein aggregates in desminopathies, and cellular compartments and processes known to play a role in desminopathies. Candidates of interest were sorted into the following categories, ‘intermediate filament & associated’, ‘sarcomere’, ‘heat shock related’, ‘protein quality control’, ‘mitochondria’, ‘oxidative stress’, ‘mechanical stress’, ‘neuromuscular junction’, ‘channels & transporters’, ‘myogenesis & regeneration’, ‘cell cycle’, ‘nuclear envelope’, ‘adhesion & migration & extracellular matrix’, “inflammation’, and ‘other candidates of interest’ (Table S6). Direct desmin interaction partners [2, 62, 63] that were detected were indicated with an asterisk (Table S6, column Q). Out of the group of ‘intermediate filament and intermediate filament associated proteins’, which harbors several direct desmin interaction partners, only two candidates, namely keratin 19 and syncoilin, were significantly regulated in both the heterozygous and homozygous genotypes on the protein level and additionally in the homozygous genotype on the mRNA level. Using the same criteria, cytochrome c oxidase subunit 7B, ATP synthase subunit a, mitochondrial fission regulator 1-like, succinate dehydrogenase cytochrome b small subunit, and ADP/ATP translocase 1 were present in the group of ‘mitochondrial proteins’, selenoprotein W in the group of ‘oxidative stress’, and calcium/calmodulin-dependent protein kinase type II subunit beta and calsequestrin-1 in the ‘other candidates of interest’ (Table S6). When focusing on the heterozygous genotype, which is comparable to the human disease, we detected not a single candidate that was regulated on both the protein and mRNA levels. Out of the genes whose mutations cause myofibrillar myopathies or protein aggregation myopathies [64, 65] (Table S6, column R), only two candidates, namely Dnajb6 and Bag3, were up-regulated on the protein level in both the heterozygous and homozygous phenotypes. When looking for proteins that were reported to be enriched in protein aggregates in skeletal muscle tissue from desminopathy patients [66] (Table S6, column S), Mlf2 (Myeloid leukemia factor 2) was up-regulated on the protein level in both the heterozygous and homozygous R405W desmin knock-in mice, and Psmb4, Xin2, and Dcxr only in the homozygous phenotype.

## Discussion

### R405W desmin causes a myofibrillar myopathy in both hetero- and homozygous desmin knock-in mice

Extending our previous work on the cardiotoxic effects of R406W/R405W mutant desmin in humans and mice [40], we report here a comprehensive characterization of the skeletal muscle pathology in hetero- and homozygous R405W desmin knock-in mice. The immunofluorescence and ultrastructural analyses depicted the pathognomic features of myofibrillar myopathies [38] comprising desmin-positive protein aggregates, degenerative changes of the myofibrillar apparatus, an increased autophagic buildup, and mitochondrial abnormalities. These classical alterations were present in both the heterozygous genotype and the homozygous R405W desmin knock-in mice. The former are the genetic equivalent of the autosomal-dominant human R406W desminopathy [2, 27, 40–45] and the latter are a surrogate for the very rare autosomal-recessive human desminopathies with maintained mutant desmin protein expression [67]. Notable is the fact that neither hetero-nor homozygous animals displayed morphological signs of protein aggregation pathology in standard H&E and Gömöri trichrome stains and that muscle weakness was only present in homozygous R405W desmin knock-in mice, which reach their age limit at 3 to 4 months due to a lethal intestinal pseudo-obstruction [40]. Thus, these findings are analogous to the previously published observations in R349P desmin knock-in mice [68], and suggest that ultrastructural changes in muscle fibers, including but not limited to protein aggregation, precede and probably underlie the development of detectable loss of muscle force.

### Noxious effects of R405W desmin on the intermediate filaments and the myofibrillar system, autophagic buildup and mitochondria

Previous *in vitro* assessment of the assembly properties of R406W desmin in equimolar mixtures with wild-type desmin showed the formation of chimeric filaments with normal morphology and interspersed sections of various irregularities, while the assembly of R406W desmin alone resulted in the formation of thickened filaments, aggregates, and fibrillar sheets [40]. The *in vitro* effects of the R406W mutant desmin were mirrored *in vivo* by desmin immunostains, which depicted multiple dot-like desmin-positive protein aggregates in the skeletal muscle of hetero- and homozygous R405W desmin knock-in mice with greater abundancy in the latter (Figure S2). A cross-striated desmin staining pattern, although often attenuated, was found only in the skeletal muscle of heterozygous animals (Figure S2B’), suggesting that the formation of chimeric filaments, as seen in the *in vitro* analysis, still allows the formation of a three-dimensional but abnormal desmin cytoskeleton. Since the cross-striated desmin immunolabeling pattern was almost completely absent in the homozygous genotype (Figure S2C,C’), the structure and function of the desmin filament system appears to be virtually abolished. On the ultrastructural level these subsarcolemmal and intermyofibrillar protein aggregates displayed a complex picture with granulo-filamentous (Figure S1A), filamentous (Figure S1C,G), granular (Figure S1D), and electron-dense (Figure S1F,H) appearances and varied in size and number between individual muscle fibers, reminiscent to the granulo-filamentous inclusions present in human desminopathies [2, 38, 69]. The presence of filamentous inclusions in homozygous animals (Figure S1G) possibly represent the equivalent of the thickened filament structures formed by pure R406W desmin *in vitro* [40]. The faulty R406W/R405W mutant desmin assembly process and its impact on the structure and function of the extrasarcomeric cytoskeleton thus provides an explanation for numerous subsequent pathological lesions in desminopathies comprising i) formation of protein aggregates, ii) defects in the proper alignment and anchorage of the myofibrillar apparatus with concomitant mechanical strain-induced degenerative alterations, iii) increased autophagic build-up due to misfolded proteins and protein aggregates, and iv) defects in the distribution, morphology and function of mitochondria. The fact that these key pathological lesions are also present in skeletal muscle tissue of autosomal-dominant and -recessive human desminopathies [2, 38] as well as in hetero- and homozygous R349P desmin knock-in mice [13, 68, 70–72] underscores the importance of such pathogenetic sequence in desminopathies.

### In search for new biomarkers: Increased blood concentrations of acylcarnitines and amino acids in homozygous R405W desmin knock-in mice

Previous studies in human desminopathies as well as in desmin knock-out and in R349P desmin knock-in mice provided evidence for morphological and functional alterations of mitochondria leading to a persistent disease-contributing secondary mitochondrial pathology [47, 72–77]. In desmin knock-out mice, the mitochondrial dysfunction was associated with widespread defects in fatty acid metabolism, as indicated by a significant decrease of the expression level of the fatty acid transporter CD36 and a significant increase in multiple acylcarnitines ranging from C3 to C18 chain length in whole blood samples [47]. Analysis of the R405W desmin knock-in mice also revealed a reduction in CD36, as well as in several other enzymes related to fatty acid and acylcarnitine metabolism, as discussed further below, at both the mRNA and protein levels, in conjunction with significantly elevated levels of several acylcarnitines ranging in chain length from C2 to C18 in homozygous animals. A constellation in which all acylcarnitines are elevated resembles a multiple acyl-CoA dehydrogenase deficiency due to a deficiency of electron transfer flavoproteins or electron transfer flavoprotein-ubiquinone oxidoreductase (ETF-QO) [78, 79], which also leads to a disorder of the dehydrogenases involved in amino acid metabolism. The generally reduced beta-oxidation in this scenario is also associated with the cytoplasmic accumulation of acetyl-CoA, pyruvate and, as observed, alanine. If and to what extent these blood parameters can serve as biomarkers for secondary mitochondrial dysfunction in autosomal-recessive desminopathies requires and deserves further clinical validation.

### The R406W mutation does not induce terminal ribosomal stalling during desmin translation

Many intermediate filament proteins, including nuclear lamins as well as cytosolic vimentin, follow a co-translational assembly mode based on the interaction of two nascent polypeptides translated by adjacent ribosomes [80]. It has been postulated that ribosomal pausing might coordinate this type of co-translational assembly [81]. While translational pausing may serve a physiological function in co-translational folding and trafficking, abnormally-long ribosomal stalls leading to ribosomal collisions may signal for ribosomal splitting (translation abortion), degradation of the unfinished nascent protein, mRNA decay, and stress response activation [82]. In this context, the proteomic analysis of soleus muscle from R405W desmin knock-in mice indicated a 2-fold increase in the levels of Ufc1 (ubiquitin-fold modifier-conjugating enzyme 1) in hetero- and homozygous mice and a halving of Ufsp2 (Ufm1-specific protease 2, involved in the conjugation and removal of Ufm1) in homozygous mice. Ufm1 is a ubiquitin-like protein modification conjugated to ER-localized ribosomes upon ribosome stalling, and has been implicated in the activation of ER-phagy and in the degradation of elongation-stalled nascent proteins associated with the Sec61 translocon [83–86]. This finding suggested that mutant desmin expression could potentially affect the cellular translation system and promote ribosomal stalling. To address if translation of the desmin transcript itself is affected, we used a dual fluorescence reporter assay, which demonstrated no incidence of abortive translation during the synthesis of either wild-type or R406W mutant desmin proteins. Therefore, it is plausible that the observed changes on the UFMylation machinery are due to the cytotoxic effects of the mutant desmin protein rather than an intrinsic problem with mutant desmin translation.

### Transcriptomic and proteomic datasets highlight changes in mitochondrial function and protein quality control in three-month-old R405W desminopathy mice

To assess disease-relevant changes, we used soleus muscle tissue from three-month-old heterozygous and homozygous R405W desmin knock-in mice and wild-type littermates to perform RNA sequencing and quantitative mass spectrometry-based proteomics. In the heterozygous genotype and much more pronounced in the homozygous genotype mRNAs were up- and down-regulated in approximately equal proportions. However, only the homozygous genotype clearly separated in the principal component analysis. On the protein level, dysregulation was also more prominent in the homozygous genotype, but markedly more proteins were up-than down-regulated, and all three genotypes were clearly separated in the principal component analysis.

A very prominent group of dysregulated candidates comprised mitochondrial proteins. Of particular interest was the down-regulation of the mitochondrial proton gradient-dependent carriers Slc25a3 (phosphate carrier), Slc25a4 (ADP/ATP translocase 1), and Slc25a12 (calcium-binding carrier Aralar1, electrogenic aspartate/glutamate antiporter), all of which have recently been described to be down-regulated in mitochondria purified from R349P desminopathy myotubes and/or in soleus muscle tissue from R349P desmin knock-in mice ([73] and supplementary tables therein). Down-regulation of these proteins provided an explanation for the detection of a reduced proton leak in mitochondria from cultivated R349P desminopathy myotubes by means of high-resolution respirometry [73]. Reduction of these highly essential carriers affects the normal mitochondrial ATP production [87] and very likely triggers a damaging cascade comprising electron transport chain overload and associated reactive oxygen species production [88] and mtDNA damage and deletions [89–91], ultimately resulting in functionally and structurally compromised mitochondria. Accordingly, our ultrastructural analysis showed enlarged mitochondria with widened cristae in the soleus muscle of R405W (Figure 3) and R349P [72] desminopathy mice. Furthermore, the down-regulation of Mrpl33 (mitochondrial 39S ribosomal protein L33, large ribosomal subunit protein bL33m) is of interest as it is a component of the mitochondrial ribosome and its knock-down was reported to cause a decrease in ATP production and an increase in reactive oxygen species [92, 93]. Notably, a homozygous mutation in *SLC25A3* has been identified as cause of reduced mitochondrial ATP production in muscle, leading to mitochondrial phosphate carrier deficiency with lactic acidosis, hypertrophic cardiomyopathy, and muscle hypotonia in humans [94]. Moreover, a homozygous mutation in *SLC25A4* has been found to cause myopathy and cardiomyopathy with multiple mtDNA deletions in skeletal muscle tissue [95].

With regard to the increased blood concentrations of multiple acylcarnitines, the down-regulation of fatty acid metabolism-related proteins is noteworthy. In particular, we detected reduced amounts of the fatty acid translocase CD36 (glycoprotein IIIb, leukocyte differentiation antigen CD36), Slc27a1 (long-chain fatty acid transport protein 1), Ces2c (acylcarnitine hydrolase), and the mitochondrial proteins arachidonate metabolism enzyme Mgst3 (glutathione S-transferase 3), Slc25a20 (carnitine/acylcarnitine carrier), Acadsb (short/branched chain specific acyl-CoA dehydrogenase) and Acadm (medium-chain specific acyl-CoA dehydrogenase). The carrier Slc25a20 is involved in the import of fatty acids of different chain lengths in the form of acylcarnitines into the mitochondrial matrix [96, 97]. The importance of this carrier is highlighted by the observation that mutations in *SLC25A20* cause carnitine-acylcarnitine translocase deficiency associated with an abnormal acylcarnitine profile and skeletal muscle damage [59, 98]. The down-regulation of Slc25a20, Acadsb and Acadm provides a mechanistic link to the elevated blood acylcarnitine levels in homozygous mice. Together with previous studies on secondary mitochondrial dysfunction in desminopathies [72–74, 77], our new results provide a more detailed insight into the mutant desmin-induced down-regulation of mitochondria-related key candidates promoting metabolic dysregulation in skeletal muscle tissue.

Given the abundance of pathological protein aggregates in both hetero- and homozygous R405W desmin knock-in mice, changes in the regulation of candidates related to protein quality control could be anticipated. Indeed, in addition to several E3 ubiquitin-protein processing proteins (Klhl21, Klhl38, Maea, March7, Rmnd5a, Sh3rf2, Trim35, Uchl1, Ufc1), our analysis depicted increased levels of autophagy-related (Sqstm1/p62, Atg101, Ulk1) and proteasome-related proteins (Gan, Usp2), but no overexpression of small heat shock proteins (HspB1/hsp27, HspB5/αB-crystallin), desmin and filamin-C, all of which have previously been described to be enriched in protein aggregates [66]. Out of the known list of myofibrillar myopathy-related genes [64], only the protein quality control related candidates Dnajb6 and Bag3 were significantly up-regulated in soleus muscle from both hetero- and homozygous R405W desmin mice at three months of age. Since our analysis also depicted markedly augmented levels of the transcription factors Mlf1 and Mlf2 (myeloid leukemia factors 1 and 2), which have been reported to bind to Dnajb6 (Mlf1 and Mlf2) [99, 100] and Bag3 (Mlf2) [99], the upregulation of this trio of proteins is of particular interest in the early disease stages. Notably, Mlf2, an interaction partner of Mlf1 [99], has been shown be enriched in protein aggregates in human desminopathies [66] thus highlighting a novel and conceivable disease link between Mlf1 and Mlf2 related transcriptional control and Dnajb6 and Bag3 mediated protein quality control.

### Effects of the expression of R405W mutant desmin on desmin binding partners, intermediate filament proteins and the myofibrillar damage marker Xin

When addressing possible effects of the R405W mutant desmin on the expression levels of known direct desmin interaction partners ((Table S6, column Q), [2, 62, 63]), solely the expression of the intermediate filament protein Sync was increased in hetero- and homozygous mice, whereas vimentin, plectin, nesprin-3, spectrin alpha chain 1, ankyrin-1, Hspb1/Hsp27, HspB5/Cryab/αB-crystallin, myomesin-1, myotubularin, calpain-1, calpain-3, and caspase-6 remained unchanged. Lamin-B2, synemin, and Stim1, were up- and nestin down-regulated, however, only in the homozygous genotype. Out of the group of intermediate filament proteins, keratin-19 was found to be down-regulated in the soleus muscle from hetero- and homozygous R405W desmin knock-in mice. This is of interest, as the absence of Krt19 in mice has been demonstrated to cause skeletal myopathy with mitochondrial and sarcolemmal reorganization [101]. In relation to the presence of marked myofibrillar damage on the ultrastructural level, our analysis also showed a significant up-regulation of Xin/Xirp2 on the protein level in both R405W desmin genotypes. Xin along with filamin-C has been described as sensitive marker for small and larger areas of myofibrillar lesions due to a rapid enrichment of both proteins to segments of sarcomeric damage at the level of Z-discs [102–105].

## Conclusions

Taken together, our study demonstrates that the expression of R405W mutant desmin leads to a myofibrillar myopathy in hetero- and homozygous R405W desmin knock-in mice. The results from heterozygous mice further highlight the notion that the presence of desmin-positive protein aggregates is not necessarily associated with muscle weakness. In line with previous results from the analysis of desmin knock-out mice, measurement of blood acylcarnitine levels appears to be of potential clinical interest in the context of recessive desminopathies. Out of the known desmin binding partners, only the expression of syncoilin was increased. With regard to myofibrillar damage, the lesion marker Xin was up-regulated in R405W desmin knock-in mice. Our data delineate a pathogenetic link between the myeloid leukemia factors 1 and 2 and the myofibrillar myopathy- and protein quality control-related proteins Dnajb6 and Bag3 in the context of desminopathies. Our main finding is that extensive secondary mitochondrial damage and proton gradient-dependent carrier-related dysfunction are important early pathogenetic steps, and therefore potential pre-clinical therapeutic targets, in R405W/R406W and R349P/R350P desminopathies. Integrated morphological, transcriptomic and proteomic data analysis is a powerful approach to identify and validate early stages of disease in desminopathies and other forms of myofibrillar myopathies.

## Acknowledgements

The authors thank the following persons and institutions: Harald Herrmann, Institute of Neuropathology, University Hospital Erlangen, Friedrich-Alexander University Erlangen-Nürnberg, Erlangen, Germany, for provision of desmin expression plasmids and R405 and 405W desmin-specific antibodies and discussion of results; Alain Schmitt, Electron Microscopy Platform, Cochin Institute, Paris, France, for providing electron microscopic images; the iGenSeq core facility at Institut du Cerveau, Paris, for RNA sequencing work; the Bioinformatics and Biostatistics Core Facility, Paris Epigenetics and Cell Fate Center, for sharing their analysis workflows; and the plateforme d’hébergement et d’expérimentation animale Buffon for mouse husbandry.

## Ethical guidelines statement

Mice were handled in accordance with European Union guidelines and French regulations for animal experimentation, and the investigations were approved by the University Paris Diderot local committee (authorization number CEB-16-2016 / 2016041216476300).

## Online supplementary information

**Supplementary Figure 1.**
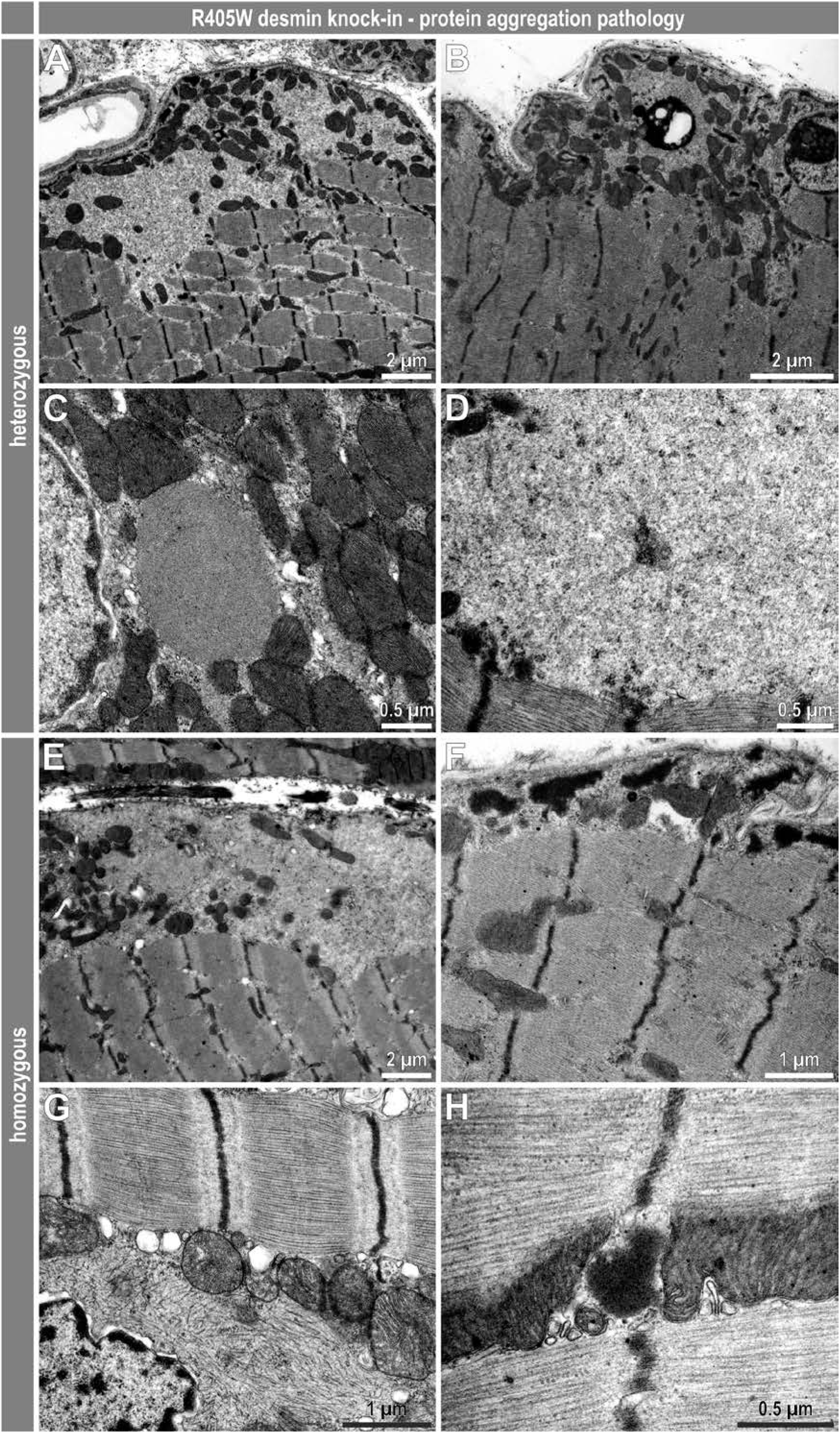
Protein aggregation pathology in skeletal muscle of heterozygous and homozygous R405W desmin knock-in mice. (**A, B**) Subsarcolemmal protein aggregates and mitochondrial accumulation in soleus muscle of heterozygous animals. Note the additional presence of a large autophagic vacuole in (B). (**C**) Cytoplasmic body adjacent to a myonucleus and surrounded by mitochondria. (**D**) Intermyofibrillar protein aggregate composed of predominantly unstructured, granular material. (**E**) Subsarcolemmal protein aggregate and mitochondrial accumulation in soleus muscle of a homozygous animal. (**F**) Subsarcolemmal protein aggregates containing electron dense material. (**G**) Filamentous protein aggregate in direct vicinity to a myonucleus, mitochondria, and a myofibril. (**H**) Electron dense protein aggregate in the intermyofibrillar space at the level of two adjacent Z-discs.

**Supplementary Figure 2.**
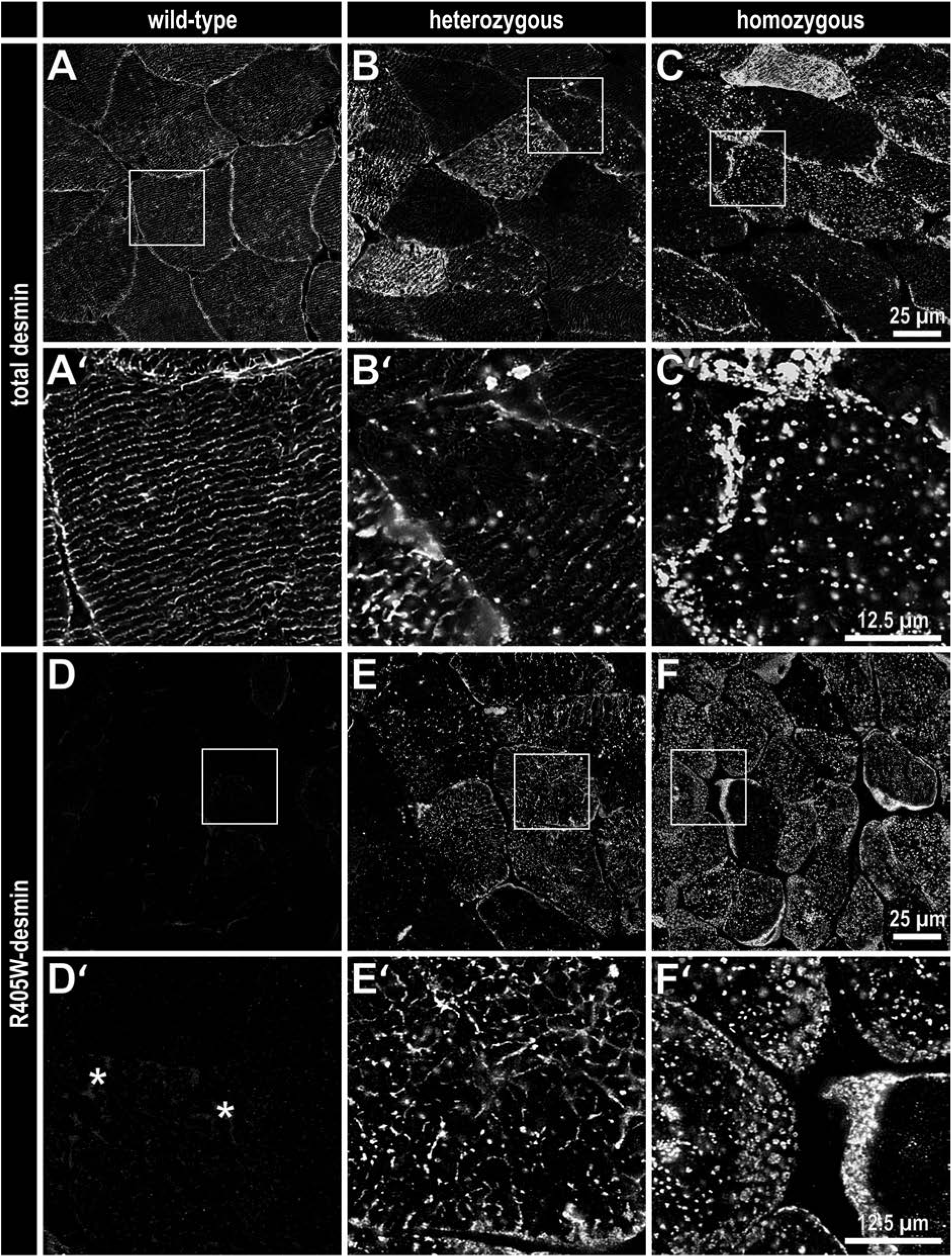
Desmin-positive protein aggregation in skeletal muscle of heterozygous and homozygous R405W desmin knock-in mice. (**A-C**) Desmin immunostaining using a desmin antibody detecting both wild-type and mutant desmin (total desmin) of transverse sections of soleus muscle from heterozygous and homozygous R405W desmin knock-in mice and wild-type littermates. Note the attenuation of the normal desmin cross-striation pattern as well as the presence of small roundish desmin-positive aggregates in heterozygous and more pronounced in homozygous muscle. (**A’-C’**) Areas with a higher magnification. (**D-F**) Analogous immunostaining using a 405W-specific desmin antibody, which visualizes small R405W desmin aggregates in the soleus muscle of heterozygous and homozygous mice. (**D’-F’**) Areas with a higher magnification. Asterisks, low-intensity autofluorescence in the subsarcolemmal region of the wild-type soleus muscle. Images were deconvolved using Huygens Essential (Scientific Volume Imaging).

[The online supplementary tables will be made available to the public upon final publication of the manuscript.]

**Supplementary Table 1. Results of the transcriptome analysis.** RNA sequencing data set from soleus muscle from heterozygous and homozygous R405W desminopathy mice and wild-type littermates.

**Supplementary Table 2. Subset of transcriptomic analysis results.** Subset of protein-coding RNAs (p<0.01, >2-fold regulation) in homozygous and wild-type soleus muscle as shown in the volcano plot (Figure 5D). mRNAs that showed an up- and down-regulation, respectively, shared between the heterozygous and homozygous genotypes are in bold orange or blue.

**Supplementary Table 3. Results of the proteome analysis.** Data set from proteotypic peptide-based quantitative proteomic analysis using soleus muscle tissue derived from hetero- and homozygous R405W desminopathy mice and wild-type littermates.

**Supplementary Table 4. Proteomics Flame analysis.** Functional enrichment analysis of the 638 significantly regulated proteins (Figure 6B, Table S3) using the Flame software (https://pavlopoulos-lab-services.org/shiny/app/flame). Enrichment results from multiple runs are presented in the ‘Combination’ tab or in separated tabs for each tool.

**Supplementary Table 5. Correlation between transcriptome and proteome analysis results.** Correlation analysis of the combined transcriptomic and proteomic datasets with a significance level of q<0.05 set for the mRNA and corresponding protein list entries resulted in four different groups of regulation; color coding as in the dot plot visualization (Figure 7).

**Supplementary Table 6. Detailed comparison of the transcriptome and proteome datasets.** Comparison, as described in the Results section, of the 4,230 entries from the combined transcriptomic and proteomic datasets. Candidates of interest were sorted into the indicated categories. Below these categories, all remaining candidates from that dataset are listed in alphabetical order.

## References

1. Lazarides E. Intermediate filaments: a chemically heterogeneous, developmentally regulated class of proteins. Annu Rev Biochem 1982;51:219–50.

2. Clemen CS, Herrmann H, Strelkov SV, Schröder R. Desminopathies: pathology and mechanisms. Acta Neuropathol 2013;125:47–75.

3. Konieczny P, Fuchs P, Reipert S, Kunz WS, Zeöld A, Fischer I, et al. Myofiber integrity depends on desmin network targeting to Z-disks and costameres via distinct plectin isoforms. J Cell Biol 2008;181:667–81.

4. O’Neill A, Williams MW, Resneck WG, Milner DJ, Capetanaki Y, Bloch RJ. Sarcolemmal organization in skeletal muscle lacking desmin: evidence for cytokeratins associated with the membrane skeleton at costameres. Mol Biol Cell 2002;13:2347–59.

5. Tidball JG. Desmin at myotendinous junctions. Exp Cell Res 1992;199:206–12.

6. Kartenbeck J, Franke WW, Moser JG, Stoffels U. Specific attachment of desmin filaments to desmosomal plaques in cardiac myocytes. EMBO J 1983;2:735–42.

7. Lapouge K, Fontao L, Champliaud MF, Jaunin F, Frias MA, Favre B, et al. New insights into the molecular basis of desmoplakin- and desmin-related cardiomyopathies. Journal of cell science 2006;119:4974–85.

8. Can T, Faas L, Ashford DA, Dowle A, Thomas J, O’Toole P, et al. Proteomic analysis of laser capture microscopy purified myotendinous junction regions from muscle sections. Proteome science 2014;12:25.

9. Eiber N, Frob F, Schowalter M, Thiel C, Clemen CS, Schröder R, et al. Lack of Desmin in Mice Causes Structural and Functional Disorders of Neuromuscular Junctions. Frontiers in molecular neuroscience 2020;13:567084.

10. Capetanaki Y, Bloch RJ, Kouloumenta A, Mavroidis M, Psarras S. Muscle intermediate filaments and their links to membranes and membranous organelles. Exp Cell Res 2007;313:2063–76.

11. Dayal AA, Medvedeva NV, Nekrasova TM, Duhalin SD, Surin AK, Minin AA. Desmin Interacts Directly with Mitochondria. Int J Mol Sci 2020;21:

12. Reipert S, Steinbock F, Fischer I, Bittner RE, Zeold A, Wiche G. Association of mitochondria with plectin and desmin intermediate filaments in striated muscle. Exp Cell Res 1999;252:479–91.

13. Diermeier S, Buttgereit A, Schürmann S, Winter L, Xu H, Murphy RM, et al. Preaged remodeling of myofibrillar cytoarchitecture in skeletal muscle expressing R349P mutant desmin. Neurobiol Aging 2017;58:77–87.

14. Ralston E, Lu Z, Biscocho N, Soumaka E, Mavroidis M, Prats C, et al. Blood vessels and desmin control the positioning of nuclei in skeletal muscle fibers. J Cell Physiol 2006;209:874–82.

15. Heffler J, Shah PP, Robison P, Phyo S, Veliz K, Uchida K, et al. A Balance Between Intermediate Filaments and Microtubules Maintains Nuclear Architecture in the Cardiomyocyte. Circ Res 2020;126:e10–e26.

16. Lieber RL, Roberts TJ, Blemker SS, Lee SSM, Herzog W. Skeletal muscle mechanics, energetics and plasticity. Journal of neuroengineering and rehabilitation 2017;14:108.

17. Palmisano MG, Bremner SN, Hornberger TA, Meyer GA, Domenighetti AA, Shah SB, et al. Skeletal muscle intermediate filaments form a stress-transmitting and stress-signaling network. J Cell Sci 2015;128:219–24.

18. Hakibilen C, Delort F, Daher MT, Joanne P, Cabet E, Cardoso O, et al. Desmin Modulates Muscle Cell Adhesion and Migration. Frontiers in cell and developmental biology 2022;10:783724.

19. Agnetti G, Herrmann H, Cohen S. New roles for desmin in the maintenance of muscle homeostasis. FEBS J 2022;289:2755–70.

20. van Spaendonck-Zwarts KY, van der Kooi AJ, van den Berg MP, Ippel EF, Boven LG, Yee WC, et al. Recurrent and founder mutations in the Netherlands: the cardiac phenotype of DES founder mutations p.S13F and p.N342D. Neth Heart J 2012;20:219–28.

21. Brodehl A, Gaertner-Rommel A, Milting H. Molecular insights into cardiomyopathies associated with desmin (DES) mutations. Biophysical reviews 2018;10:983–1006.

22. Carmignac V, Sharma S, Arbogast S, Fischer D, Serreri C, Serria M, et al. A homozygous desmin deletion causes an Emery-Dreifuss like recessive myopathy with desmin depletion. Neuromuscul Disord 2009;19:600.

23. Henderson M, De Waele L, Hudson J, Eagle M, Sewry C, Marsh J, et al. Recessive desmin-null muscular dystrophy with central nuclei and mitochondrial abnormalities. Acta Neuropathol 2013;125:917–9.

24. McLaughlin HM, Kelly MA, Hawley PP, Darras BT, Funke B, Picker J. Compound heterozygosity of predicted loss-of-function DES variants in a family with recessive desminopathy. BMC Med Genet 2013;14:68.

25. Durmus H, Ayhan O, Cirak S, Deymeer F, Parman Y, Franke A, et al. Neuromuscular endplate pathology in recessive desminopathies: Lessons from man and mice. Neurology 2016;87:799–805.

26. Batonnet-Pichon S, Behin A, Cabet E, Delort F, Vicart P, Lilienbaum A. Myofibrillar Myopathies: New Perspectives from Animal Models to Potential Therapeutic Approaches. Journal of neuromuscular diseases 2017;4:1–15.

27. Dagvadorj A, Olive M, Urtizberea JA, Halle M, Shatunov A, Bonnemann C, et al. A series of West European patients with severe cardiac and skeletal myopathy associated with a de novo R406W mutation in desmin. J Neurol 2004;251:143–9.

28. Fichna JP, Karolczak J, Potulska-Chromik A, Miszta P, Berdynski M, Sikorska A, et al. Two desmin gene mutations associated with myofibrillar myopathies in Polish families. PloS one 2014;9:e115470.

29. Fidzianska A, Kotowicz J, Sadowska M, Goudeau B, Walczak E, Vicart P, et al. A novel desmin R355P mutation causes cardiac and skeletal myopathy. Neuromuscul Disord 2005;15:525–31.

30. Goldfarb LG, Park KY, Cervenakova L, Gorokhova S, Lee HS, Vasconcelos O, et al. Missense mutations in desmin associated with familial cardiac and skeletal myopathy. Nat Genet 1998;19:402–3.

31. Mavroidis M, Panagopoulou P, Kostavasili I, Weisleder N, Capetanaki Y. A missense mutation in desmin tail domain linked to human dilated cardiomyopathy promotes cleavage of the head domain and abolishes its Z-disc localization. FASEB J 2008;22:3318–27.

32. Pica EC, Kathirvel P, Pramono ZA, Lai PS, Yee WC. Characterization of a novel S13F desmin mutation associated with desmin myopathy and heart block in a Chinese family. Neuromuscul Disord 2008;18:178–82.

33. Arbustini E, Morbini P, Grasso M, Fasani R, Verga L, Bellini O, et al. Restrictive cardiomyopathy, atrioventricular block and mild to subclinical myopathy in patients with desmin-immunoreactive material deposits. J Am Coll Cardiol 1998;31:645–53.

34. Cetin N, Balci-Hayta B, Gundesli H, Korkusuz P, Purali N, Talim B, et al. A novel desmin mutation leading to autosomal recessive limb-girdle muscular dystrophy: distinct histopathological outcomes compared with desminopathies. J Med Genet 2013;50:437–43.

35. Munoz-Marmol AM, Strasser G, Isamat M, Coulombe PA, Yang Y, Roca X, et al. A dysfunctional desmin mutation in a patient with severe generalized myopathy. Proc Natl Acad Sci USA 1998;95:11312–7.

36. Pinol-Ripoll G, Shatunov A, Cabello A, Larrode P, de la Puerta I, Pelegrin J, et al. Severe infantile-onset cardiomyopathy associated with a homozygous deletion in desmin. Neuromuscul Disord 2009;19:418–22.

37. Riley LG, Waddell LB, Ghaoui R, Evesson FJ, Cummings BB, Bryen SJ, et al. Recessive DES cardio/myopathy without myofibrillar aggregates: intronic splice variant silences one allele leaving only missense L190P-desmin. Eur J Hum Genet 2019;27:1267–73.

38. Schröder R, Schoser B. Myofibrillar myopathies: a clinical and myopathological guide. Brain Pathol 2009;19:483–92.

39. Schröder R. Protein aggregate myopathies: the many faces of an expanding disease group. Acta Neuropathol 2013;125:1–2.

40. Herrmann H, Cabet E, Chevalier NR, Moosmann J, Schultheis D, Haas J, et al. Dual functional states of R406W-desmin assembly complexes cause cardiomyopathy with severe intercalated disc derangement in humans and in knock-in mice. Circulation 2020;142:2155–71.

41. Arbustini E, Pasotti M, Pilotto A, Pellegrini C, Grasso M, Previtali S, et al. Desmin accumulation restrictive cardiomyopathy and atrioventricular block associated with desmin gene defects. Eur J Heart Fail 2006;8:477–83.

42. Dalakas MC, Park KY, Semino-Mora C, Lee HS, Sivakumar K, Goldfarb LG. Desmin myopathy, a skeletal myopathy with cardiomyopathy caused by mutations in the desmin gene. N Engl J Med 2000;342:770–80.

43. Olive M, Goldfarb L, Moreno D, Laforet E, Dagvadorj A, Sambuughin N, et al. Desmin-related myopathy: clinical, electrophysiological, radiological, neuropathological and genetic studies. J Neurol Sci 2004;219:125–37.

44. Park KY, Dalakas MC, Semino-Mora C, Lee HS, Litvak S, Takeda K, et al. Sporadic cardiac and skeletal myopathy caused by a de novo desmin mutation. Clin Genet 2000;57:423–9.

45. Wahbi K, Behin A, Charron P, Dunand M, Richard P, Meune C, et al. High cardiovascular morbidity and mortality in myofibrillar myopathies due to DES gene mutations: a 10-year longitudinal study. Neuromuscul Disord 2012;22:211–8.

46. Mill L, Aust O, Ackermann JA, Burger P, Pascual M, Palumbo-Zerr K, et al. SYNTA: A novel approach for deep learning-based image analysis in muscle histopathology using photo-realistic synthetic data. 2022. p. arXiv:2207.14650.

47. Elsnicova B, Hornikova D, Tibenska V, Kolar D, Tlapakova T, Schmid B, et al. Desmin Knock-Out Cardiomyopathy: A Heart on the Verge of Metabolic Crisis. Int J Mol Sci 2022;23:12020.

48. Demichev V, Messner CB, Vernardis SI, Lilley KS, Ralser M. DIA-NN: neural networks and interference correction enable deep proteome coverage in high throughput. Nat Methods 2020;17:41–4.

49. Deutsch EW, Bandeira N, Perez-Riverol Y, Sharma V, Carver JJ, Mendoza L, et al. The ProteomeXchange consortium at 10 years: 2023 update. Nucleic Acids Res 2023;51:D1539–D48.

50. Perez-Riverol Y, Bai J, Bandla C, Garcia-Seisdedos D, Hewapathirana S, Kamatchinathan S, et al. The PRIDE database resources in 2022: a hub for mass spectrometry-based proteomics evidences. Nucleic Acids Res 2022;50:D543–D52.

51. Zhang X, Jonassen I. RASflow: an RNA-Seq analysis workflow with Snakemake. BMC Bioinformatics 2020;21:110.

52. Dobin A, Davis CA, Schlesinger F, Drenkow J, Zaleski C, Jha S, et al. STAR: ultrafast universal RNA-seq aligner. Bioinformatics 2013;29:15–21.

53. Liao Y, Smyth GK, Shi W. The R package Rsubread is easier, faster, cheaper and better for alignment and quantification of RNA sequencing reads. Nucleic Acids Res 2019;47:e47.

54. Love MI, Huber W, Anders S. Moderated estimation of fold change and dispersion for RNA-seq data with DESeq2. Genome biology 2014;15:550.

55. Andersen CL, Jensen JL, Orntoft TF. Normalization of real-time quantitative reverse transcription-PCR data: a model-based variance estimation approach to identify genes suited for normalization, applied to bladder and colon cancer data sets. Cancer Res 2004;64:5245–50.

56. Ran FA, Hsu PD, Wright J, Agarwala V, Scott DA, Zhang F. Genome engineering using the CRISPR-Cas9 system. Nat Protoc 2013;8:2281–308.

57. Lackner DH, Carré A, Guzzardo PM, Banning C, Mangena R, Henley T, et al. A generic strategy for CRISPR-Cas9-mediated gene tagging. Nature Communications 2015;6:10237.

58. Upper D. The unsuccessful self-treatment of a case of “writer’s block“. J Appl Behav Anal 1974;7:497.

59. Dambrova M, Makrecka-Kuka M, Kuka J, Vilskersts R, Nordberg D, Attwood MM, et al. Acylcarnitines: Nomenclature, Biomarkers, Therapeutic Potential, Drug Targets, and Clinical Trials. Pharmacol Rev 2022;74:506–51.

60. Juszkiewicz S, Hegde RS. Initiation of Quality Control during Poly(A) Translation Requires Site-Specific Ribosome Ubiquitination. Mol Cell 2017;65:743–50 e4.

61. Sundaramoorthy E, Leonard M, Mak R, Liao J, Fulzele A, Bennett EJ. ZNF598 and RACK1 Regulate Mammalian Ribosome-Associated Quality Control Function by Mediating Regulatory 40S Ribosomal Ubiquitylation. Mol Cell 2017;65:751–60 e4.

62. Hernandez DA, Bennett CM, Dunina-Barkovskaya L, Wedig T, Capetanaki Y, Herrmann H, et al. Nebulette is a powerful cytolinker organizing desmin and actin in mouse hearts. Mol Biol Cell 2016;27:3869–82.

63. Zhang H, Bryson VG, Wang C, Li T, Kerr JP, Wilson R, et al. Desmin interacts with STIM1 and coordinates Ca2+ signaling in skeletal muscle. JCI insight 2021;6:

64. Olivé M, Winter L, Fürst DO, Schröder R, group Ews. 246th ENMC International Workshop: Protein aggregate myopathies 24-26 May 2019, Hoofddorp, The Netherlands. Neuromuscul Disord 2021;31:158–66.

65. Weihl CC, Topf A, Bengoechea R, Duff J, Charlton R, Garcia SK, et al. Loss of function variants in DNAJB4 cause a myopathy with early respiratory failure. Acta Neuropathol 2023;145:127–43.

66. Maerkens A, Kley RA, Olive M, Theis V, van der Ven PF, Reimann J, et al. Differential proteomic analysis of abnormal intramyoplasmic aggregates in desminopathy. J Proteomics 2013;90:14–27.

67. Ruppert T, Heckmann MB, Rapti K, Schultheis D, Jungmann A, Katus HA, et al. AAV-mediated cardiac gene transfer of wild-type desmin in mouse models for recessive desminopathies. Gene Ther 2020;27:516–24.

68. Clemen CS, Stöckigt F, Strucksberg KH, Chevessier F, Winter L, Schütz J, et al. The toxic effect of R350P mutant desmin in striated muscle of man and mouse. Acta Neuropathol 2015;129:297–315.

69. Claeys KG, van der Ven PF, Behin A, Stojkovic T, Eymard B, Dubourg O, et al. Differential involvement of sarcomeric proteins in myofibrillar myopathies: a morphological and immunohistochemical study. Acta Neuropathol 2009;117:293–307.

70. Diermeier S, Iberl J, Vetter K, Haug M, Pollmann C, Reischl B, et al. Early signs of architectural and biomechanical failure in isolated myofibers and immortalized myoblasts from desmin-mutant knock-in mice. Scientific reports 2017;7:1391.

71. Winter L, Unger A, Berwanger C, Spörrer M, Türk M, Chevessier F, et al. Imbalances in protein homeostasis caused by mutant desmin. Neuropathol Appl Neurobiol 2019;45:476–94.

72. Winter L, Wittig I, Peeva V, Eggers B, Heidler J, Chevessier F, et al. Mutant desmin substantially perturbs mitochondrial morphology, function and maintenance in skeletal muscle tissue. Acta Neuropathol 2016;132:453–73.

73. Berwanger C, Terres D, Pesta D, Eggers B, Marcus K, Wittig I, et al. Immortalised murine R349P desmin knock-in myotubes exhibit a reduced proton leak and decreased ADP/ATP translocase levels in purified mitochondria. Eur J Cell Biol 2024;103:151399.

74. Hovhannisyan Y, Li Z, Callon D, Suspene R, Batoumeni V, Canette A, et al. Critical contribution of mitochondria in the development of cardiomyopathy linked to desmin mutation. Stem cell research & therapy 2024;15:10.

75. Lindén M, Li Z, Paulin D, Gotow T, Leterrier JF. Effects of desmin gene knockout on mice heart mitochondria. J Bioenerg Biomembr 2001;33:333–41.

76. Schröder R, Goudeau B, Simon MC, Fischer D, Eggermann T, Clemen CS, et al. On noxious desmin: functional effects of a novel heterozygous desmin insertion mutation on the extrasarcomeric desmin cytoskeleton and mitochondria. Hum Mol Genet 2003;12:657–69.

77. Smolina N, Khudiakov A, Knyazeva A, Zlotina A, Sukhareva K, Kondratov K, et al. Desmin mutations result in mitochondrial dysfunction regardless of their aggregation properties. Biochimica et biophysica acta Molecular basis of disease 2020;1866:165745.

78. Niederwieser A, Steinmann B, Exner U, Neuheiser F, Redweik U, Wang M, et al. Multiple acyl-Co A dehydrogenation deficiency (MADD) in a boy with nonketotic hypoglycemia, hepatomegaly, muscle hypotonia and cardiomyopathy. Detection of N-isovalerylglutamic acid and its monoamide. Helv Paediatr Acta 1983;38:9–26.

79. Cornelius N, Frerman FE, Corydon TJ, Palmfeldt J, Bross P, Gregersen N, et al. Molecular mechanisms of riboflavin responsiveness in patients with ETF-QO variations and multiple acyl-CoA dehydrogenation deficiency. Hum Mol Genet 2012;21:3435–48.

80. Bertolini M, Fenzl K, Kats I, Wruck F, Tippmann F, Schmitt J, et al. Interactions between nascent proteins translated by adjacent ribosomes drive homomer assembly. 2021;371:57–64.

81. Panasenko OO, Somasekharan SP, Villanyi Z, Zagatti M, Bezrukov F, Rashpa R, et al. Co-translational assembly of proteasome subunits in NOT1-containing assemblysomes. Nature Structural & Molecular Biology 2019;26:110–20.

82. Eisenack TJ, Trentini DB. Ending a bad start: Triggers and mechanisms of co-translational protein degradation. Frontiers in molecular biosciences 2022;9:1089825.

83. Wang L, Xu Y, Rogers H, Saidi L, Noguchi CT, Li H, et al. UFMylation of RPL26 links translocation-associated quality control to endoplasmic reticulum protein homeostasis. Cell Res 2020;30:5–20.

84. Stephani M, Picchianti L, Gajic A, Beveridge R, Skarwan E, Sanchez de Medina Hernandez V, et al. A cross-kingdom conserved ER-phagy receptor maintains endoplasmic reticulum homeostasis during stress. Elife 2020;9:1–105.

85. Scavone F, Gumbin Samantha C, Da Rosa Paul A, Kopito Ron R. RPL26/uL24 UFMylation is essential for ribosome-associated quality control at the endoplasmic reticulum. 2023;120:e2220340120.

86. Liang JR, Lingeman E, Luong T, Ahmed S, Muhar M, Nguyen T, et al. A Genome-wide ER-phagy Screen Highlights Key Roles of Mitochondrial Metabolism and ER-Resident UFMylation. Cell 2020;180:1160–77.e20.

87. Mishra G, Coyne LP, Chen XJ. Adenine nucleotide carrier protein dysfunction in human disease. IUBMB life 2023;75:911–25.

88. Wallace DC, Fan W, Procaccio V. Mitochondrial energetics and therapeutics. Annual review of pathology 2010;5:297–348.

89. Beckman KB, Ames BN. The free radical theory of aging matures. Physiol Rev 1998;78:547–81.

90. Dos Santos JM, de Oliveira DS, Moreli ML, Benite-Ribeiro SA. The role of mitochondrial DNA damage at skeletal muscle oxidative stress on the development of type 2 diabetes. Mol Cell Biochem 2018;449:251–5.

91. Wiesner RJ, Zsurka G, Kunz WS. Mitochondrial DNA damage and the aging process: facts and imaginations. Free Radic Res 2006;40:1284–94.

92. Liu L, Luo C, Luo Y, Chen L, Liu Y, Wang Y, et al. MRPL33 and its splicing regulator hnRNPK are required for mitochondria function and implicated in tumor progression. Oncogene 2018;37:86–94.

93. Brown A, Amunts A, Bai XC, Sugimoto Y, Edwards PC, Murshudov G, et al. Structure of the large ribosomal subunit from human mitochondria. Science 2014;346:718–22.

94. Mayr JA, Merkel O, Kohlwein SD, Gebhardt BR, Bohles H, Fotschl U, et al. Mitochondrial phosphate-carrier deficiency: a novel disorder of oxidative phosphorylation. Am J Hum Genet 2007;80:478–84.

95. Echaniz-Laguna A, Chassagne M, Ceresuela J, Rouvet I, Padet S, Acquaviva C, et al. Complete loss of expression of the ANT1 gene causing cardiomyopathy and myopathy. Journal of medical genetics 2012;49:146–50.

96. De Lucas JR, Indiveri C, Tonazzi A, Perez P, Giangregorio N, Iacobazzi V, et al. Functional characterization of residues within the carnitine/acylcarnitine translocase RX2PANAAXF distinct motif. Mol Membr Biol 2008;25:152–63.

97. Perez P, Martinez O, Romero B, Olivas I, Pedregosa AM, Palmieri F, et al. Functional analysis of mutations in the human carnitine/acylcarnitine translocase in Aspergillus nidulans. Fungal Genet Biol 2003;39:211–20.

98. Iacobazzi V, Invernizzi F, Baratta S, Pons R, Chung W, Garavaglia B, et al. Molecular and functional analysis of SLC25A20 mutations causing carnitine-acylcarnitine translocase deficiency. Hum Mutat 2004;24:312–20.

99. Taipale M, Tucker G, Peng J, Krykbaeva I, Lin ZY, Larsen B, et al. A quantitative chaperone interaction network reveals the architecture of cellular protein homeostasis pathways. Cell 2014;158:434–48.

100. Blandin G, Marchand S, Charton K, Daniele N, Gicquel E, Boucheteil JB, et al. A human skeletal muscle interactome centered on proteins involved in muscular dystrophies: LGMD interactome. Skeletal muscle 2013;3:3.

101. Stone MR, O’Neill A, Lovering RM, Strong J, Resneck WG, Reed PW, et al. Absence of keratin 19 in mice causes skeletal myopathy with mitochondrial and sarcolemmal reorganization. Journal of cell science 2007;120:3999–4008.

102. Chevessier F, Schuld J, Orfanos Z, Plank AC, Wolf L, Maerkens A, et al. Myofibrillar instability exacerbated by acute exercise in filaminopathy. Hum Mol Genet 2015;24:7207–20.

103. Schuld J, Orfanos Z, Chevessier F, Eggers B, Heil L, Uszkoreit J, et al. Homozygous expression of the myofibrillar myopathy-associated p.W2710X filamin C variant reveals major pathomechanisms of sarcomeric lesion formation. Acta neuropathologica communications 2020;8:154.

104. Nilsson MI, Nissar AA, Al-Sajee D, Tarnopolsky MA, Parise G, Lach B, et al. Xin is a marker of skeletal muscle damage severity in myopathies. Am J Pathol 2013;183:1703–9.

105. Shatunov A, Olive M, Odgerel Z, Stadelmann-Nessler C, Irlbacher K, van Landeghem F, et al. In-frame deletion in the seventh immunoglobulin-like repeat of filamin C in a family with myofibrillar myopathy. Eur J Hum Genet 2009;17:656–63.

106. Lin YJ, Huang LH, Huang CT. Enhancement of heterologous gene expression in Flammulina velutipes using polycistronic vectors containing a viral 2A cleavage sequence. PLoS One 2013;8:e59099.

107. Oliveros JC. Venny. An interactive tool for comparing lists with Venn’s diagrams. 2015. https://bioinfogp.cnb.csic.es/tools/venny/index.html.

